# An orphan gene is essential for efficient sperm entry into eggs in *Drosophila melanogaster*

**DOI:** 10.1101/2024.08.08.607187

**Authors:** Sara Y. Guay, Prajal H. Patel, Jonathon M. Thomalla, Kerry L. McDermott, Jillian M. O’Toole, Sarah E. Arnold, Sarah J. Obrycki, Mariana F. Wolfner, Geoffrey D. Findlay

**Affiliations:** Department of Biology, College of the Holy Cross, Worcester, MA 01610; Department of Molecular Biology and Genetics, Cornell University, Ithaca, NY 14853

**Keywords:** orphan gene, *Drosophila*, sperm, reproduction, molecular evolution, fertilization

## Abstract

While spermatogenesis has been extensively characterized in the *Drosophila melanogaster* model system, very little is known about the genes required for fly sperm entry into eggs. We identified a lineage-specific gene, which we named *katherine johnson* (*kj*), that is required for efficient fertilization. Males that do not express *kj* produce and transfer sperm that are stored normally in females, but sperm from these males enter eggs with severely reduced efficiency. Using a tagged transgenic rescue construct, we observed that the KJ protein localizes around the edge of the nucleus at various stages of spermatogenesis but is undetectable in mature sperm. These data suggest that *kj* exerts an effect on sperm development, the loss of which results in reduced fertilization ability. Interestingly, KJ protein lacks detectable sequence similarity to any other known protein, suggesting that *kj* could be a lineage-specific orphan gene. While previous bioinformatic analyses indicated that *kj* was restricted to the *melanogaster* group of *Drosophila*, we identified putative orthologs with conserved synteny, male-biased expression, and predicted protein features across the genus, as well as likely instances of gene loss in some lineages. Thus, *kj* was likely present in the *Drosophila* common ancestor and subsequently evolved an essential role in fertility in *D. melanogaster*. Our results demonstrate a new aspect of male reproduction that has been shaped by a lineage-specific gene and provide a molecular foothold for further investigating the mechanism of sperm entry into eggs in *Drosophila*.

**Article Summary:** How fruit fly sperm enter eggs is poorly understood. Here, we identify a gene required for efficient fertilization. Sperm from males lacking this gene’s function cannot enter eggs. The gene appears to act during sperm production, rather than in mature sperm. Interestingly, the gene is undetectable outside of genus *Drosophila*, and its encoded protein shows no discernable similarity to other proteins. This study provides insights into sperm-egg interactions and illustrates how lineage-specific genes can impact important aspects of reproduction.

## Introduction

In many animal species, fertilization is a complex, yet essential, process that requires the successful production of sperm, the transfer to and storage of sperm within females, the entry of a sperm into an egg cell, and the correct unpackaging and use of paternal chromatin. The first part of this process, spermatogenesis, has been well characterized in a variety of systems, including *Drosophila (Fabian and Brill 2012)*, and has broadly similar features across metazoans (White-Cooper et al. 2009). What happens after sperm leave the male, but before development begins, is an active area of study, about which less is known. Upon transfer to females, sperm must navigate through the reproductive tract to reach specialized site(s) at which they can be stored (Wolfner et al. 2023). In mammals, sperm storage typically involves binding to specialized regions of the oviduct epithelium (Suarez 2008), while in insects, specialized sperm storage organs are used (Pitnick et al. 1999). Stored sperm must then be released at a rate appropriate to fertilize oocytes when the latter are ovulated (Bloch Qazi et al. 2003; Manier et al. 2010). Upon release, sperm must find the egg and then fertilize it. In many taxa, including mammals and marine invertebrates, initial interactions between sperm and egg include the sperm’s acrosome reaction (Okabe 2016), which facilitates the fusion of the sperm and egg plasma membranes and allows the contents of the sperm nucleus to enter the egg (Deneke and Pauli 2021; Elofsson et al. 2024). In *Drosophila* and some fish species, however, a sperm cell gains access to the egg through a cone-shaped projection in the eggshell called the micropyle (Horne-Badovinac 2020). How *Drosophila* sperm locate the micropyle is unknown, as is the mechanism through which the entire *Drosophila* sperm cell passes through the egg plasma membrane. The identification of a fly mutant in which sperm were unable to enter eggs (Perotti et al. 2001) suggested the possibility that specific gene products could be responsible for either of these steps, but that fly line is no longer available, and its affected gene was never identified molecularly. After a fly sperm enters an egg, the sperm plasma membrane breaks down, releasing a lysosome (the former acrosome), the nucleus, and centrioles. This membrane breakdown is mediated by a sperm transmembrane protein, Sneaky, and is required for the subsequent unpackaging of the paternal genome (Fitch and Wakimoto 1998; Wilson et al. 2006). After the paternal genome is released, additional male- and female-derived proteins are required for proper chromatin decondensation and use (Loppin et al. 2001; Loppin, Bonnefoy, et al. 2005; Sakai et al. 2009; Tirmarche et al. 2016; Yamaki et al. 2016; Dubruille et al. 2023); mutations in the genes encoding these proteins lead to paternal- or maternal-effect lethality, respectively. Although *Drosophila* genetics has enabled the identification of many of these components (Loppin et al. 2015), our understanding of the processes between spermatogenesis and the onset of development remains incomplete.

While many aspects of spermatogenesis are conserved, sperm are also among the fastest evolving cell types, likely due to sexual selection (Pitnick et al. 2009; Ramm et al. 2014). Across genus *Drosophila*, species produce different numbers of sperm per differentiated germline stem cell (Schärer et al. 2008), sperm length is highly variable (Lüpold et al. 2016), and males of some species produce multiple types of sperm (Alpern et al. 2019). Correspondingly, females of different species have evolved diverse structures for and patterns of sperm storage (Pitnick et al. 1999). These observations suggest a role for lineage-specific evolution in shaping sperm traits. Such evolution could occur through changes to the coding sequences (Wilburn and Swanson 2016) and/or expression patterns (VanKuren and Long 2018) existing genes. Sperm traits could also evolve through the formation of lineage-specific genes through processes such as gene duplication, gene fusion, horizontal gene transfer or *de novo* gene birth (Long et al. 2013).

Numerous lineage-specific genes have evolved important roles in *Drosophila* spermatogenesis. For example, arising through recent duplication and subsequent regulatory evolution, the *nsr* gene regulates the expression of several Y-linked genes required for sperm individualization and axoneme formation (Ding et al. 2010). The *ms(3)K81* gene arose in the *melanogaster* group of *Drosophila* through retrotransposition and is required for protecting the telomeres of paternal chromatin during fertilization (Loppin, Lepetit, et al. 2005; Dubruille et al. 2010).

Lineage-specific duplications of the highly conserved *Arp2* gene, which promotes actin filament nucleation, have evolved testis-specific expression in the *montium* and *obscura* groups of *Drosophila*, and insertion of these paralogs into *D. melanogaster* disrupts spermatogenesis (Stromberg et al. 2023). VanKuren and Long (VanKuren and Long 2018) demonstrated that the duplication of a gene that was likely expressed in both male and female germlines in the ancestor of *D. melanogaster* gave rise to paralogs that evolved either testis- or ovary-specific expression, with the male-specific gene, *Apollo*, now being required for spermatid individualization. In addition to these lineage-specific genes that arose via duplication-based processes, we previously identified three genes that appeared to be restricted to the *Drosophila* genus, lacked detectable homology to any other protein, and were essential for robust male fertility (Gubala et al. 2017; Lange et al. 2021; Rivard et al. 2021). For example, *goddard* encodes a protein that localizes to developing sperm axonemes and is required for proper spermatid individualization (Lange et al. 2021), while *atlas* encodes a protein that localizes to spermatid nuclei and appears to transiently bind DNA during the process of nuclear condensation (Rivard et al. 2021). Because these genes appear restricted to the *Drosophila* genus and encode proteins with no detectable homology to other proteins, we initially described them as putatively *de novo* evolved.

*De novo* gene evolution occurs when mutations transform a previously non-coding segment of the genome into a protein-coding gene (Van Oss and Carvunis 2019; L. Zhao et al. 2024). To establish a gene as *de novo* evolved, the syntenic region should be identified in outgroup species and confirmed to be non-genic. This is most feasible for *de novo* genes that are evolutionarily young, so the highest-confidence *de novo* genes are those that are found in only one or a few species and for which closely related outgroup species have genome sequence data available (Levine et al. 2006; Begun et al. 2007; Carvunis et al. 2012; Zhao et al. 2014; Zhang et al. 2019; Vakirlis et al. 2020; Vakirlis et al. 2022). Older genes that appear lineage-restricted and lack detectable homology, but for which a syntenic, non-coding region is not identifiable in outgroup species, have historically been called putative *de novo* genes (McLysaght and Hurst 2016; Van Oss and Carvunis 2019), the term that we applied to genes such as *goddard* and *atlas* (Gubala et al. 2017; Lange et al. 2021; Rivard et al. 2021). As the *de novo* gene field has matured, however, researchers have recognized that issues such as the limited sensitivity of sequence-based homology searches and the breakdown of synteny in progressively more diverged genomes can cause distant homologs of putative *de novo* genes to be missed (Weisman et al. 2020; L. Zhao et al. 2024). Thus, such genes might now be referred to more cautiously as “orphans” (Tautz and Domazet-Lošo 2011; Q. Zhao et al. 2024). This broader term describes lineage-specific genes that lack detectable homologs outside of a particular clade for any reason (e.g., *de novo* origin, divergence beyond recognition, gene loss in outgroup species, horizontal gene transfer, or genome assembly issues).

One potential advance in distinguishing *de novo* genes from other types of orphans is the use of whole-genome alignments (Peng and Zhao 2024). This approach facilitates the identification of the syntenic region in diverged species, which in turn limits the search space for sequence homology searches, improving their sensitivity. Peng and Zhao (2024) used this approach to identify hundreds of likely *de novo* genes in *D. melanogaster* and, equally importantly, to distinguish other orphans that either had a different origin or for which the origin could not be definitively determined. Despite this significant advance, both early (Wagstaff and Begun 2005; Findlay et al. 2009) and more recent (Gubala et al. 2017; Rivard et al. 2021) experience with cross-species reproductive gene annotation in *Drosophila* suggests that manual annotation of individual genes can sometimes identify orthologs that were undetected by high-throughput bioinformatic analyses.

Here, we investigated the male reproductive function and molecular evolution of the *D. melanogaster* gene *CG43167*, which we have named *katherine johnson* (*kj*). This gene was identified in two bioinformatic screens (Heames et al. 2020; Peng and Zhao 2024) as likely *de novo* evolved and restricted to the *melanogaster* group of *Drosophila*. We show here that knockdown or knockout of *kj* results in a severe reduction in male fertility. Knockout males produce sperm that are stored at normal levels in females’ seminal receptacles, but the sperm enter eggs at much reduced rates. Because the KJ protein is detectable in various stages of spermatogenesis, but not in mature sperm, we suggest that *kj* exerts its effect during sperm development, and that in its absence, the ability of sperm to fertilize eggs is significantly impaired. Across the *melanogaster* group of *Drosophila* species, *kj* has maintained a male-biased pattern of expression but shows an elevated rate of sequence evolution. By analyzing gene synteny, expression patterns, and predicted protein features, we identified putative orthologs in outgroup *Drosophila* species, as well as lineages in which the gene is undetectable. These data suggest *kj* was present at the base of the *Drosophila* genus, but might have become expendable in certain lineages as spermatogenic processes diverged. The likely presence of *kj* in a more ancient ancestor makes it harder to determine whether the gene evolved *de novo*, so we consider *kj* to be an orphan gene. Overall, our study provides a potential foothold from which to further our understanding of *Drosophila* fertilization, highlights a critical reproductive role in *D. melanogaster* for an orphan gene, and illustrates a challenge of large-scale bioinformatic identification of *de novo* genes.

## Methods

### Drosophila stocks and experiments

Please see the Reagents Table for a full list of fly strains used in this study. Unless otherwise noted, *in vivo* experiments in *Drosophila* were performed at 25°C using standard molasses media consisting of agar (6.5 g/L), brewers yeast (23.5 g/L), cornmeal (60 g/L), molasses (60 mL/L), acid mix (4 mL/L; propionic and phosphoric acids), and tegosept (0.13%; antifungal agent).

### Genetic ablation of CG43167

We first constructed a TRiP-style RNAi line (Ni et al. 2011) targeting *CG43167* expression and used RT-PCR to assess the degree of knockdown. The oligos used for creating the pValium20 plasmid and for RT-PCR are provided in Fig. S1. Fertility of small groups of knockdown and control male flies was assessed as previously described (Rivard et al. 2021).

We used the co-CRISPR method as previously described (Ge et al. 2016; Lange et al. 2021; Rivard et al. 2021) to engineer a complete deletion of *CG43167*. Guide RNA sequences used to target the gene and PCR primers used to verify the deletion are provided in Fig. S2. Flies carrying a deletion allele (Δ*kj*) were crossed into the *w^1118^* background and balanced over CyO. We generated trans-heterozygotes with no functional copies of *kj* using Bloomington Stock Center deficiency line #9717, with genotype *w^1118^*; Df(2L)BSC243/CyO.

Unless otherwise stated, heterozygous control flies used in experiments were generated by crossing the Δ*kj* line to *w*^1118^; we refer to these controls as Δ*kj*/+.

### Cloning and transformation of tagged kj rescue constructs

C-terminally tagged *kj:HA* rescue construct and N-terminally tagged *HA:kj* rescue constructs were generated using Gibson Assembly (Gibson et al. 2009). The *kj* coding sequence and putative upstream and downstream regulatory sequences were amplified from Canton S genomic DNA (prepared using Gentra Puregene Cell Kit, Qiagen) using Q5 High Fidelity Polymerase (NEB). Primers used for making all constructs are listed in the Reagents Table. The 3x-HA tag was similarly amplified using pTWH plasmids (T. Murphy, *Drosophila* Genomics Resource Center plasmids 1100 and 1076). Amplified DNA fragments were then assembled into a XbaI/AscI-linearized *w*+ attB plasmid (a gift of Jeff Sekelsky, Addgene plasmid 30326).

Assembled constructs were integrated into the attP docking site of PBac{*y*^+^-attP-3B}VK00037 (Bloomington *Drosophila* Stock Center stock #24872) using PhiC31 integrase (Rainbow Transgenics).

### Fertility assays

Male fertility of *kj* nulls, flies carrying rescue constructs, and controls was assessed using matings between single unmated males of each genotype and single unmated Canton S females. Males and females were collected and isolated for a period of 72-96 hours prior to mating. During this period, females were reared in yeasted vials to encourage egg production. Each pair mating was then allowed to proceed for 72 hours before the parents were removed from the vial. Fertility was determined by counting pupal cases on the side of vials 10 days after the initial crossing. Twenty matings were set up for each male genotype; vials with any dead parents or atypical bacterial growth at the end of the mating period were excluded from analysis.

### Sperm counts

We crossed the *Mst35Bb*-GFP (“protamine-GFP”) marker of mature sperm nuclei (Manier et al. 2010) into the *kj* null background and used it to quantify levels of sperm in the seminal vesicles of sexually mature, unmated males (3-5 days old), in the bursae of females 30 minutes after the start of mating (ASM), in the female seminal receptacle 2 hours ASM, and in the female seminal receptacle 4 days ASM. Matings, dissections, imaging and counting were performed as previously described (Gubala et al. 2017). Experimenters were blinded to the male genotype while counting sperm. Two-sample *t*-tests with unequal variances were used to compare sperm levels.

### Egg-production and egg-to-pupae viability assay

We measured the amount of egg-laying, the rate at which eggs developed into pupae, and the total progeny production of Canton S females mated singly to either a *kj* null male or a heterozygous control (Δ*kj*/+) using standard assays largely as previously described (Ravi Ram and Wolfner 2007; LaFlamme et al. 2012; Findlay et al. 2014). However, because the effects of *kj* knockout were large and consistent across days, we modified these procedures by: measuring egg-laying over four days (with one vial per female per day) instead of 10; analyzing pooled data across all four days of the assay (after observing that each individual day showed the same pattern); and using two-sample *t*-tests with unequal variances to compare knockout and control genotypes for each set of pooled data.

### Sperm entry into eggs and early embryonic development

We recombined the *Dj*-GFP sperm tail marker (Santel et al. 1997) into the Δ*kj* null background. For experiments examining sperm entry and early development, fly strains were maintained on yeast-glucose-agar media (Hu et al. 2020).

### Embryo collection

All embryo collections were performed at room temperature. For each embryo collection cage, approximately 30 2-7 day-old males were mated overnight to approximately 40 3-6 day-old Canton S females. Embryos were collected on grape juice agar plates (2.15% agar, 49% grape juice, 0.5% propionic acid solution (86.3% acid/water mix)) with yeast paste smeared on top. To assess embryo development, plates with embryos were collected after approximately 18 hours. For Dj-GFP detection, embryos were pre-collected for 1 hour to allow flies to lay any retained eggs. Then, fresh grape juice plates with yeast paste were replaced in 1 hour intervals.

### Sperm tail detection using Dj-GFP

Embryos from 1 hour collection plates were immediately dechorionated by treating with 50% bleach for 2 minutes. Embryos were then washed thoroughly with egg wash buffer (0.4% NaCl, 0.03% Triton-X100) and transferred to a 22x60mm coverslip prepared with a thin strip of heptane glue (stabilizes embryos lined up in a row to prevent double counting). Excess egg wash buffer was added to the slide to prevent embryo dehydration. Embryos were then imaged live on an Echo Revolve at 10X magnification to determine the proportion with detectable Dj-GFP sperm tails. For display purposes, some embryos were also fixed and imaged with confocal microscopy as described below. To ensure mating occurred, females from embryo collection cages were dissected and reproductive tracts were imaged to confirm presence of Dj-GFP sperm in the storage organs.

### Embryo development assay

Embryos collected overnight were dechorionated with 50% bleach for 2 minutes and washed thoroughly with egg wash buffer. Embryos were then fixed for 20 minutes at room temperature in 1:1 mixture of 4% paraformaldehyde in 1X PBS and heptane. Embryos were devitellinized in a 1:1 mixture of heptane and methanol by shaking vigorously for 30 seconds. Embryos were then washed three times in both pure methanol followed by 1X PBS-T (0.1% Triton-X100). To detect nuclei, embryos were stained for 20 minutes at room temperature with 10mM Hoechst 33342 diluted 1:1000 and then washed thrice with 1X PBS-T. Embryos were then mounted on 22x22mm coverslips in Aqua Polymount. Embryos were imaged on a Zeiss LSM710 confocal microscope. Images were captured using either EC-Plan Neofluar 10x/0.45 Air or Plan-Apochromat 63x/1.4 oil objectives.

### Cytology of KJ subcellular localization

We performed whole testis staining as described in Lange et al. (2021). Analysis of KJ expression in isolated cysts was performed as described in Rivard et al. (2021). We tested for KJ in mature sperm by aging male flies in single-sex vials for 10-14 days prior to dissection to allow sperm to accumulate in the seminal vesicles. Seminal vesicles were then dissected on 0.01% poly-L-lysine treated slides and pierced to release their sperm content. See the Reagents Table for details on primary and secondary antibodies. Labeled samples were imaged using a TCS SP8 X confocal microscope (Leica Microsystems). Images were captured using HC PL APO CS2 20x/0.75 ILL and HC PL APO CS2 63x/1.40 oil objectives.

Post-acquisition processing was performed using ImageJ Fiji (version 1.0).

### Sperm nuclei decondensation assay

Nuclear decondensation was performed using a modified protocol described by Tirmarche et al. (2016). Sperm were isolated from aged seminal vesicles as described above. Sperm nuclei were subsequently decondensed by pretreating sperm with 1X PBS (phosphate buffered saline) supplemented with 1% Triton X-100 for 30 minutes prior to subjecting sperm to decondensation buffer (10 mM DTT and 500 ug/mL heparin sodium salt in 1X PBS). Following treatment, slides were stained with anti-HA antibodies using the immunohistochemistry protocol described (Rivard et al. 2021).

### Molecular evolutionary analyses

We extracted the *kj* protein-coding DNA sequence and predicted amino acid sequence for *D. melanogaster* from FlyBase (Öztürk-Çolak et al. 2024). We used the protein as a query in iterative PSI-BLAST searches, which identified annotated orthologs across the *melanogaster* group of *Drosophila.* Because these orthologs varied in the quality of their annotations, we manually checked all orthologs for which genome browsers and RNA-seq data were available through the Genomics Education Partnership (thegep.org). Briefly, we BLASTed the predicted protein sequence of each PSI-BLAST hit against the corresponding species’ genome assembly, then manually examined that species’ genome in the GEP’s implementation of the UCSC Genome Browser (Rele et al. 2022). This allowed us to visualize adult male and adult female RNA-seq reads (Brown et al. 2014; Chen et al. 2014) that mapped to the region so that we could assess expression patterns. To search for orthologs outside of the *melanogaster* group, we examined the syntenic region in outgroup species (Rivard et al. 2021; Rele et al. 2022) as demarcated by three conserved genes with conserved positions relative to each other and to *kj*: *CG6614*, *CG4983* and *Vha100-5.* Any unannotated location in the syntenic region that showed adult male expression by RNA-seq was examined for potential open reading frames, and potential proteins so identified were compared to *D. melanogaster* (and other) KJ orthologs and to the full *D. melanogaster* proteome by BLASTP. We examined the predicted membrane topology of potential orthologs with DeepTMHMM (Hallgren et al. 2022). Finally, potential orthologs found in non-*melanogaster* group species were compared by BLASTP to other *Drosophila* orthologs and by BLASTP and PSI-BLAST to all known proteins in GenBank.

We examined the molecular evolution of *kj* protein-coding sequences from the *melanogaster* group as described previously (Rivard et al. 2021). In addition to those PAML-based tests of positive selection, we implemented HyPhy-based tests for recurrent (Kosakovsky Pond and Frost 2005) and episodic (Murrell et al. 2015; Wisotsky et al. 2020) positive selection as implemented in the Datamonkey 2.0 web server (Weaver et al. 2018). The sequence alignment used in these analyses was checked for recombination using GARD (Kosakovsky Pond et al. 2006), but none was detected.

## Results

### *CG43167* is required for full male fertility

*CG43167* was identified as a potential *de novo* evolved gene in two previous bioinformatic analyses (Heames et al. 2020; Peng and Zhao 2024) and shows a highly testis-biased pattern of expression (Vedelek et al. 2018). We found that expression of a short hairpin targeting *CG43167* using the *Bam-GAL4, UAS-Dicer2* driver had a marked effect on male fertility. Crude fertility assays in which seven knockdown or control males were mated with five unmated wild-type (Canton S) females for 2 days showed knockdown male fertility to be only 7-19% the level of controls. RT-PCR analysis of cDNA synthesized from controls and knockdown males showed virtually no detectable expression in knockdown males, suggesting that the transgenic line efficiently targets *CG43167* transcripts (Fig. S1). Consistent with our previous rocket-themed nomenclature for testis-expressed orphan genes (Gubala et al. 2017; Rivard et al. 2021), we named the *CG43167* gene *katherine johnson* (*kj*), after the NASA mathematician who calculated rocket orbital mechanics for the Mercury and subsequent crewed missions (Shetterly 2016).

To confirm these data and to generate a null allele for genetic analysis, we engineered a deletion of the *kj*/*CG43167* gene region using CRISPR/Cas9. The resulting deletion allele (*Δkj*) eliminated the entirety of the protein-coding and untranslated regions and thus most likely constitutes a functional null (Fig. S2). Single pair fertility assays, in which either single control males (*w^1118^)* or single *Δkj* homozygous null males were individually mated to single, wildtype, unmated females, revealed that *Δkj* null males have a fertility defect of a similar magnitude to that observed in the RNAi assay (Fig. 1). To rule out the effects of off-target mutations generated during CRISPR/Cas9 genome editing, we assessed the fertility of heterozygous males carrying a single copy of the *Δkj* allele in trans with *Df(2L)BSC243* (henceforth abbreviated as “*Df*”), a large genomic deficiency that uncovers several genes including the *kj* locus. In single pair fertility assays, *Δkj*/*Df* trans-heterozygous males showed a fertility defect equivalent to *Δkj* null males, indicating that the severe loss-of-function phenotype in *Δkj* homozygotes reflects a full loss of *kj* function (Fig. 1). To further characterize the *Δkj* allele, we determined the fertility of male flies carrying only one copy of the *Δkj* allele. Removing a single copy of the *kj* gene had no effect on male fertility, ruling out dominance by haploinsufficiency (Fig. 1). Altogether, these experiments show that the *Δkj* allele acts as a recessive null allele.

**Figure 1.**
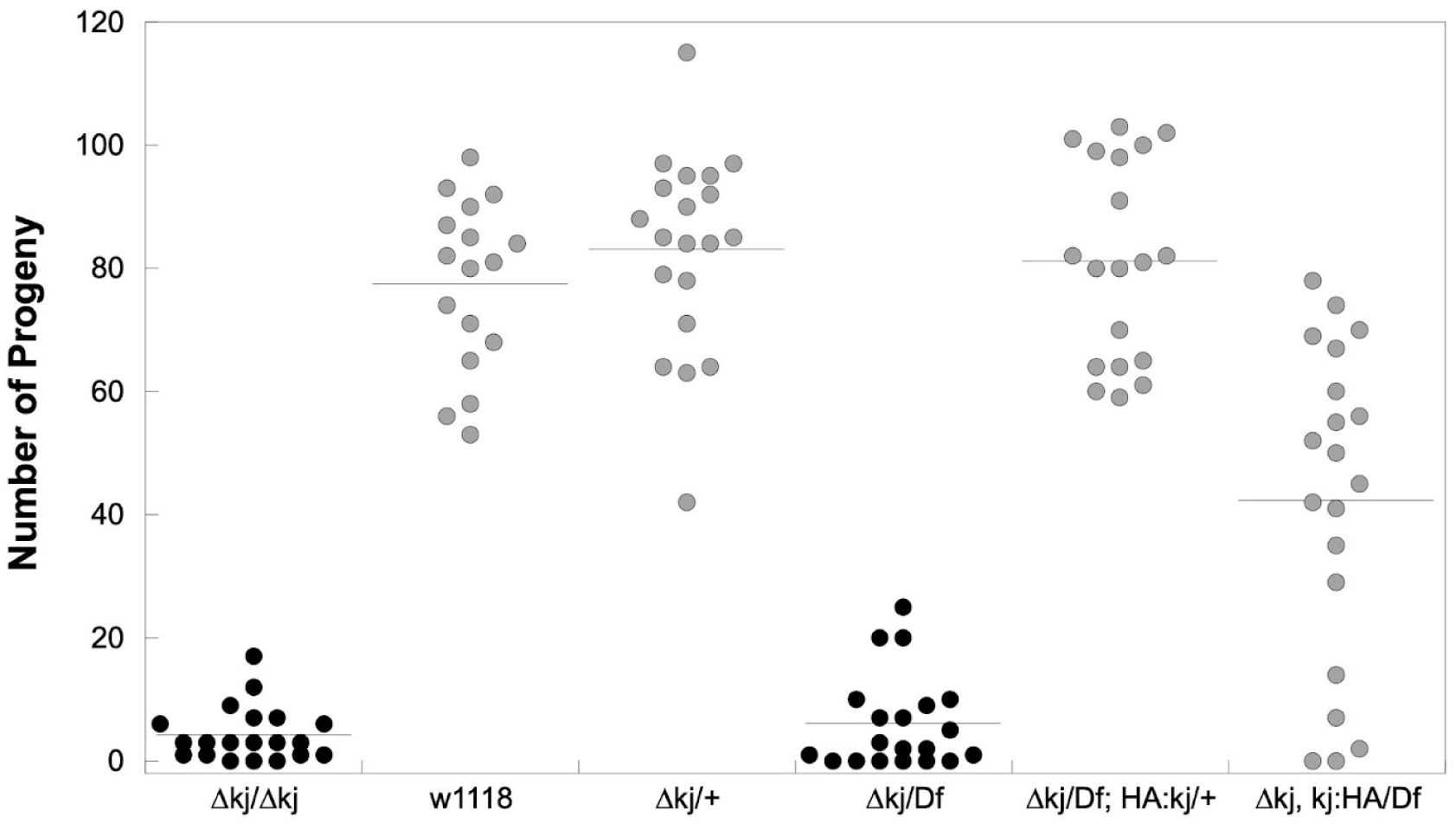
The *katherine johnson* gene (*CG43167*) is required for maximal male fertility in *D. melanogaster*. Males homozygous for a complete deletion allele (Δ*kj*) had significantly lower fertility than *w*^1118^ controls and Δ*kj*/+ heterozygotes (both *p* < 10^-13^). Δ*kj*/+ heterozygotes had no significant fertility difference from *w*^1118^ (*p* = 0.26), indicating the Δ*kj* allele is fully recessive. Trans-heterozygote males (Δ*kj*/*Df*) with no functional copies of *kj* showed no significant difference in fertility relative to Δ*kj* homozygotes (*p* = 0.37). Fertility of Δ*kj*/*Df* heterozygotes was significantly increased upon addition of either of two tagged rescue constructs, HA:kj (*p* < 10^-15^) or kj:HA (*p* < 10^-5^). The N-terminally tagged construct had significantly higher fertility than the C-terminally tagged construct (*p* < 10^-5^) and showed no significant fertility difference from *w*^1118^ controls (*p* = 0.46). Progeny number was counted as the number of pupal cases produced by females mated to males of a specific genotype. All *p*-values are from two-tailed *t*-tests with unequal variances. Horizontal lines show means. Samples sizes were *n* = 17-20 per genotype.

We confirmed that the fertility defects associated with *Δkj* are due to loss of the *kj*/*CG43167* gene by complementing the loss of function phenotype with genomic rescue constructs. We integrated the 5.4-kb *kj* locus, which contained the 583 bp *CG43167* transcript-encoding sequence along with putative upstream and downstream regulatory regions. No other annotated genes are present in this stretch of DNA. Two different constructs were produced for this analysis, differing in either the N-terminal or C-terminal location of an introduced hemagglutinin (3xHA) tag. Reintroducing either construct into *Δkj*/*Df* males restored fertility (Fig. 1). However, the degree of rescue with the C-terminally tagged protein (KJ:HA) was weaker than that of the N-terminally tagged protein (HA:KJ), which showed full fertility restoration (Fig. 1). Thus, for the remainder of the study, we focused on the N-terminally-tagged rescue construct. Collectively, these data indicate that the *kj* gene has an essential function in *Drosophila melanogaster* male fertility.

### *kj* null males produce, transfer and store sperm normally, but the sperm enter eggs inefficiently

When we examined testis morphology in *kj* null males, we observed no gross differences from control testes (Fig. S3). Furthermore, sperm with apparently normal morphology were present in the seminal vesicles (SV) of both control and mutant tracts, suggesting that spermatogenesis can proceed to completion in the absence of *kj* function. We used the *Mst35Bb-*GFP sperm head marker (Manier et al. 2010) to quantify sperm present in SVs of sexually mature, unmated males. We found a slight decrease in the number of sperm per SV in *kj* null males relative to controls (Table 1). While statistically significant, this difference was not of the same magnitude as the observed fertility difference (Fig. 1) and therefore cannot account for the observed fertility defects in Δ*kj* males.

**Table 1.**
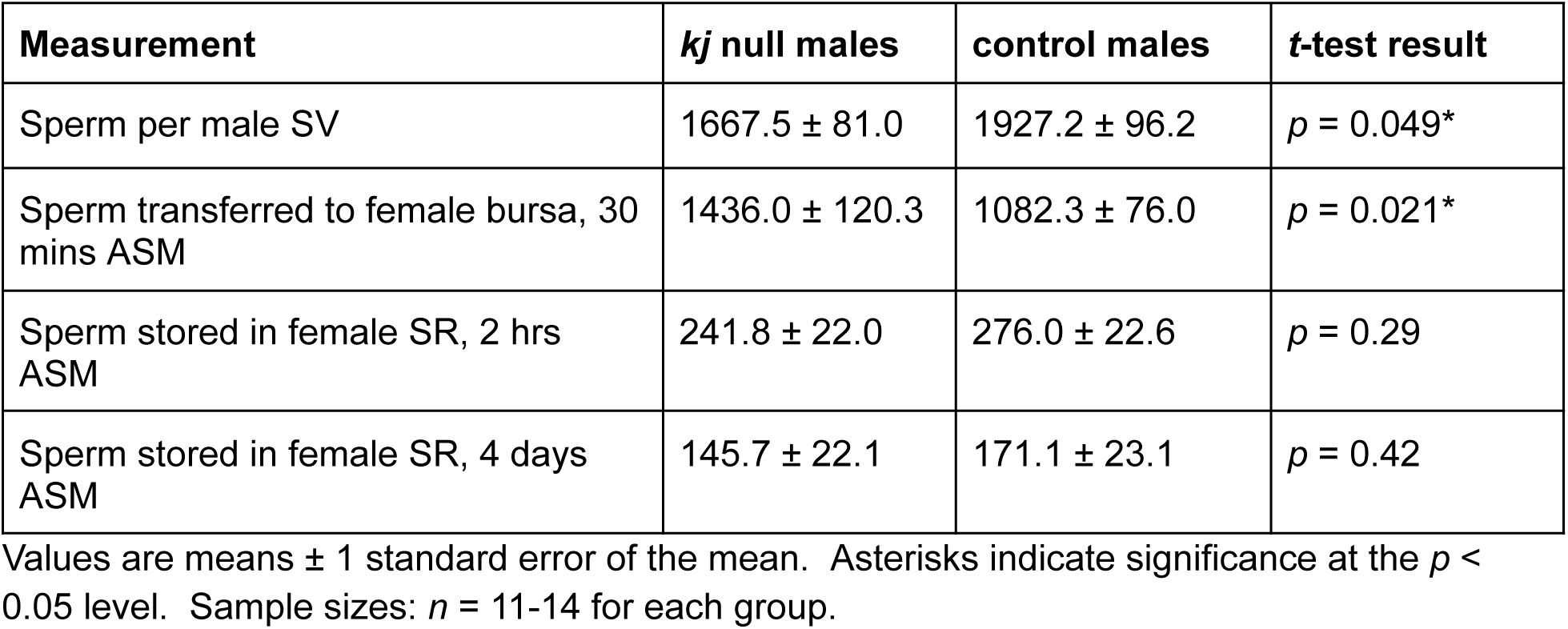
Sperm production, transfer and storage for *kj* null males and controls.

In addition to producing mature sperm, *D. melanogaster* males must also transfer sperm into females and generate functional sperm that can swim to female storage organs (Manier et al. 2010). We assessed sperm transfer by counting sperm in the female bursa (or uterus) 30 minutes after the start of mating (ASM), and observed the opposite pattern, a slight but significant increase in sperm transferred by *kj* null males (Table 1). Again, this difference was not of a comparable magnitude to the null fertility defect, nor was it in the expected direction. Thus, while *kj* null males may exhibit minor differences from controls in sperm production and sperm transfer to females, neither is likely to be the primary cause of the *kj* null fertility defect.

Since *D. melanogaster* sperm must enter specialized sperm storage organs before they can be used for fertilization, we next quantified sperm levels in the female’s primary storage organ, the seminal receptacle (SR), at two timepoints (Table 1). The level of sperm in the SR at 2 hrs ASM indicates the ability of sperm to enter storage, while sperm levels at 4 days ASM provide a readout of sperm persistence in storage and the rate of sperm release during the initial days after mating. Females mated to *kj* null males showed no significant differences in the levels of stored sperm at either time point (Table 1). Thus, sperm from *kj* null males migrate to and enter the SR normally and appear to be released from the SR at a comparable rate to sperm from heterozygous controls.

We next assessed the rates of egg laying and egg-to-pupal viability in females mated singly to either *kj* null or control males. In the four days following mating, females mated to *kj* null males laid a slightly, but not statistically significantly, lower number of eggs compared to females mated to controls (Fig. 2A). However, a much lower percentage of these eggs hatched (i.e., developed to pupae) (Fig. 2B), and accordingly, mates of *kj* nulls produced lower levels of progeny (Fig. 2C). Taken together with the sperm storage data (Table 1), these results suggest that the *kj* null fertility defect arises within a narrow, but critical, window of time between the release of sperm from storage and the onset of development.

**Figure 2.**
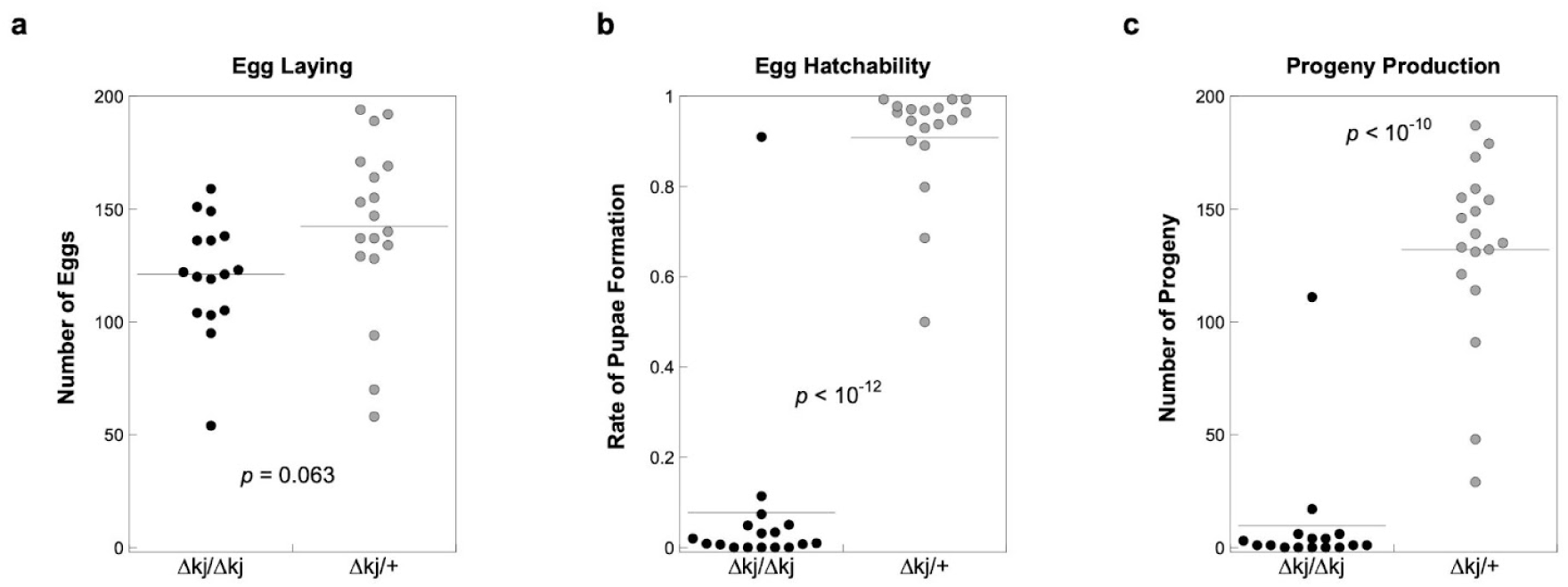
The fertility defect of *kj* null males results from an egg hatching defect. a) Egg-laying over a four-day assay by females mated to *kj* null males or heterozygous (Δ*kj*/+) controls. The groups showed no significant difference. b) The proportion of eggs from panel (a) that developed to pupae. Eggs laid by mates of *kj* null males had a significantly lower hatching rate. c) Progeny production for females mated to *kj* null males is correspondingly lower. The single high outlier for the *kj* null genotype in panels b and c might have resulted from the use of a mis-identified Δ*kj*/+ heterozygous male in the *kj* null group. In each panel, horizontal lines indicate means, and the two genotypes were compared by two-sample *t*-tests with unequal variances, with *p*-values given in the graphs.

As Δ*kj* males produced sperm that can be maintained in storage and do not hamper egg laying in females, we reasoned that the *kj* fertility defect may be due to either an inability of mutant sperm to enter eggs (Perotti et al. 2001) or a defect in a step immediately following sperm entry. Sperm with defects in the latter process fall into the category of paternal effect lethals and reflect aberrations in post-fertilizations events, such as failures in sperm plasma membrane breakdown (Wilson et al. 2006) or in the proper decondensation or initial use of the paternal chromatin inside the embryo (Loppin, Lepetit, et al. 2005; Dubruille et al. 2023).

To distinguish these possibilities, we crossed the *don juan-*GFP (*Dj*-GFP) marker (Santel et al. 1997) into the *kj* null background. This marker labels mature sperm tails and allows for the visualization of sperm entry into eggs. Canton S (wild-type) females were mated to either Δ*kj*/CyO or Δ*kj*/Δ*kj* males expressing *Dj*-GFP and allowed to lay eggs on grape juice plates in one-hour intervals. Eggs were then immediately dechorionated and imaged live by epifluorescence to assess sperm presence in the anterior end of the embryo (for examples of embryos with and without sperm, see fixed confocal images in Fig. 3A-B; example epifluorescence images used for quantification are in Fig. S4). While nearly 80% of embryos laid by females mated to heterozygous males had detectable sperm tails, *Dj-*GFP was detected in only 0.74% of embryos laid by females mated to *kj* null males (Fig. 3A-C). This significant decrease in sperm entry rate was consistent with the magnitude of the fertility differences observed above (Fig. 1, Fig. 2C), so we concluded that the inability of sperm to enter eggs efficiently is the major factor driving the *kj* null subfertility phenotype.

**Figure 3:**
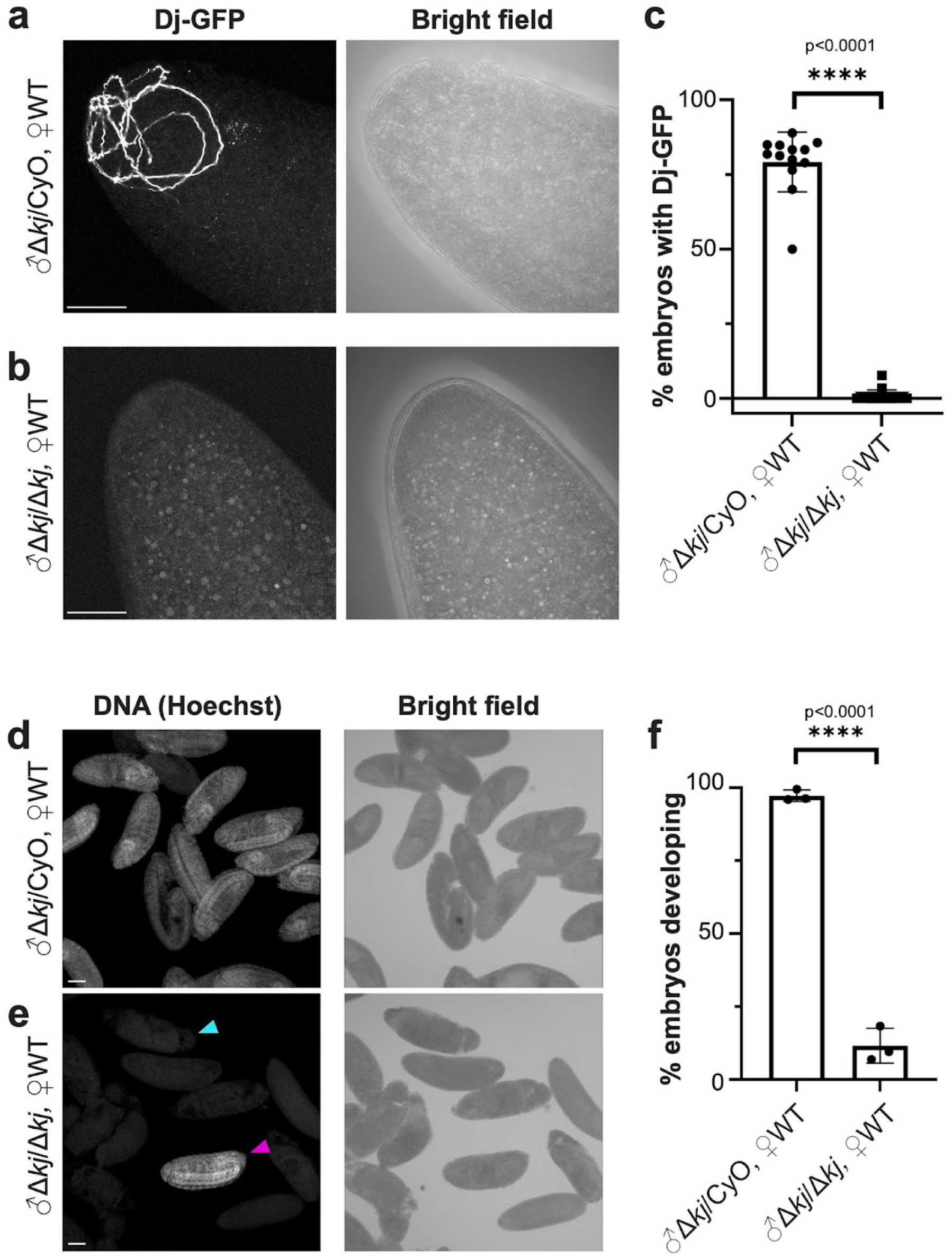
S**p**erm **from *kj* null males fertilize eggs inefficiently.** a-c) Max projection confocal images of fixed <1 hour old embryos laid by Canton S (WT) females mated to either Δ*kj*/CyO controls or Δ*kj*/Δ*kj* males expressing *Dj*-GFP (scale bars = 50μm). a) *Dj*-GFP sperm from Δ*kj*/CyO flies were frequently detected in the anterior of <1 hour old WT embryos. b) *Dj*-GFP sperm from Δ*kj*/Δ*kj* flies were rarely detected in the anterior of <1 hour old WT embryos. c) Quantification of a,b. Embryos fathered by Δ*kj*/CyO flies are positive for *Dj*-GFP 79.2% of the time (n=212 embryos), compared to 0.7% when fathered by Δ*kj*/Δ*kj* flies (n=275 embryos). d-f) Max projection confocal images of fixed, Hoechst-stained embryos collected overnight from WT females mated to either Δ*kj*/CyO or Δ*kj*/Δ*kj* males (scale bars = 100μm). d) Embryos fertilized by Δ*kj*/CyO males develop normally and reach up to Stage 16 of embryonic development during the collection period. e) When fertilized by Δ*kj*/Δ*kj* males, embryos appear to develop normally (magenta arrowhead). Unfertilized embryos deteriorate during the collection period (cyan arrowhead). f) Quantification of d,e. Embryos from females mated to Δ*kj*/CyO males appear to develop normally 97.3% of the time (n=504 embryos), compared to 11.6% of the time when mated to Δ*kj*/Δ*kj* males (n=544 embryos). ****p<0.0001, unpaired t-test, two-tailed. At least three biological replicates were performed for each experiment.

To evaluate the possibility of an additional defect in embryos successfully fertilized by Δ*kj*/Δ*kj* sperm, mated females were allowed to lay eggs onto grape juice plates for an 18-hour overnight period. Embryos were then collected and stained for DNA to allow us to assess embryonic development. Over 97% of embryos laid by females mated to control males developed normally, with a mix of developing stages up to Stage 16 present as expected (Fig. 3D, F; exact stages not quantified) (Foe et al. 1993). However, embryos laid by females mated to Δ*kj*/Δ*kj* males showed normally developing embryos only 11.6% of the time (Fig. 3E, magenta arrowhead, Fig. 3F), with similar stages present as controls. The remaining 88.4% of embryos were devoid of DNA staining and appeared to have deteriorated (Fig. 3E, cyan arrowhead), consistent with the embryos being successfully laid and activated, but not fertilized (Horner and Wolfner 2008). These experiments indicate that the few eggs that are successfully fertilized by sperm from Δ*kj*/Δ*kj* males can progress normally through embryogenesis, consistent with the outcomes of our fertility assays. Thus, *kj* expression in the male germline appears not to affect development (i.e., *kj* is not a paternal effect gene), and the *kj* null fertility defect occurs between the time of sperm exit from storage and entry into eggs.

### KJ protein localizes around the edge of the nucleus during spermatogenesis but is not detected in mature sperm

To investigate potential KJ protein functions, we used the fully functional HA:KJ rescue construct (Fig. 1) in the *kj* null background to examine the expression pattern and subcellular localization of KJ protein within male reproductive tracts. Although *kj* mutants show no major defect in sperm production, we detected HA:KJ in the testes at specific stages of spermatogenesis (Fig. 4A). In spermatocytes (pre-meiotic cells), HA:KJ was enriched around the edge of the nucleus and was observed diffusely in the cytoplasm (Fig. 4B). HA:KJ was also present in post-meiotic spermatids. In these cells, bundled nuclei synchronously proceed through a stepwise condensation process that ultimately produces the thin sperm heads found in mature sperm (Rathke et al. 2014). Round and canoe shaped nuclear bundles reflect elongating stages of spermiogenesis, while needle shaped nuclei, with their fully condensed chromosomes, characterize spermatids undergoing individualization. Analysis of spermatid cysts revealed that HA:KJ localizes transiently around the nucleus during the canoe stages before disappearing at the onset of individualization (Fig. 4C). HA:KJ showed an asymmetric localization in these cells, with enrichment along one long edge of each nucleus. This pattern is reminiscent of proteins that localize to the dense body, a structure that develops during elongation and disappears at the onset of individualization (Fabian and Brill 2012; Li et al. 2023). Consistent with the disappearance of HA:KJ from nuclei at individualization, anti-HA staining of mature sperm isolated from SVs did not detect HA:KJ around the nucleus (Fig. 4D). To investigate whether HA:KJ is no longer localized around mature sperm nuclei, or whether it became inaccessible to our antibody due to the extreme degree of nuclear condensation in mature sperm (Eren-Ghiani et al. 2015; Kaur et al. 2022), we performed the same staining after decondensing mature sperm nuclei *in vitro*. Although it was not possible to perform a positive control, the strength and shape of the DNA signal changed in response to this procedure, likely reflecting at least some decondensation. However, HA:KJ remained undetectable (Fig. S5). Overall, these data suggest that KJ plays a role in sperm development that affects later sperm function in females.

**Figure 4.**
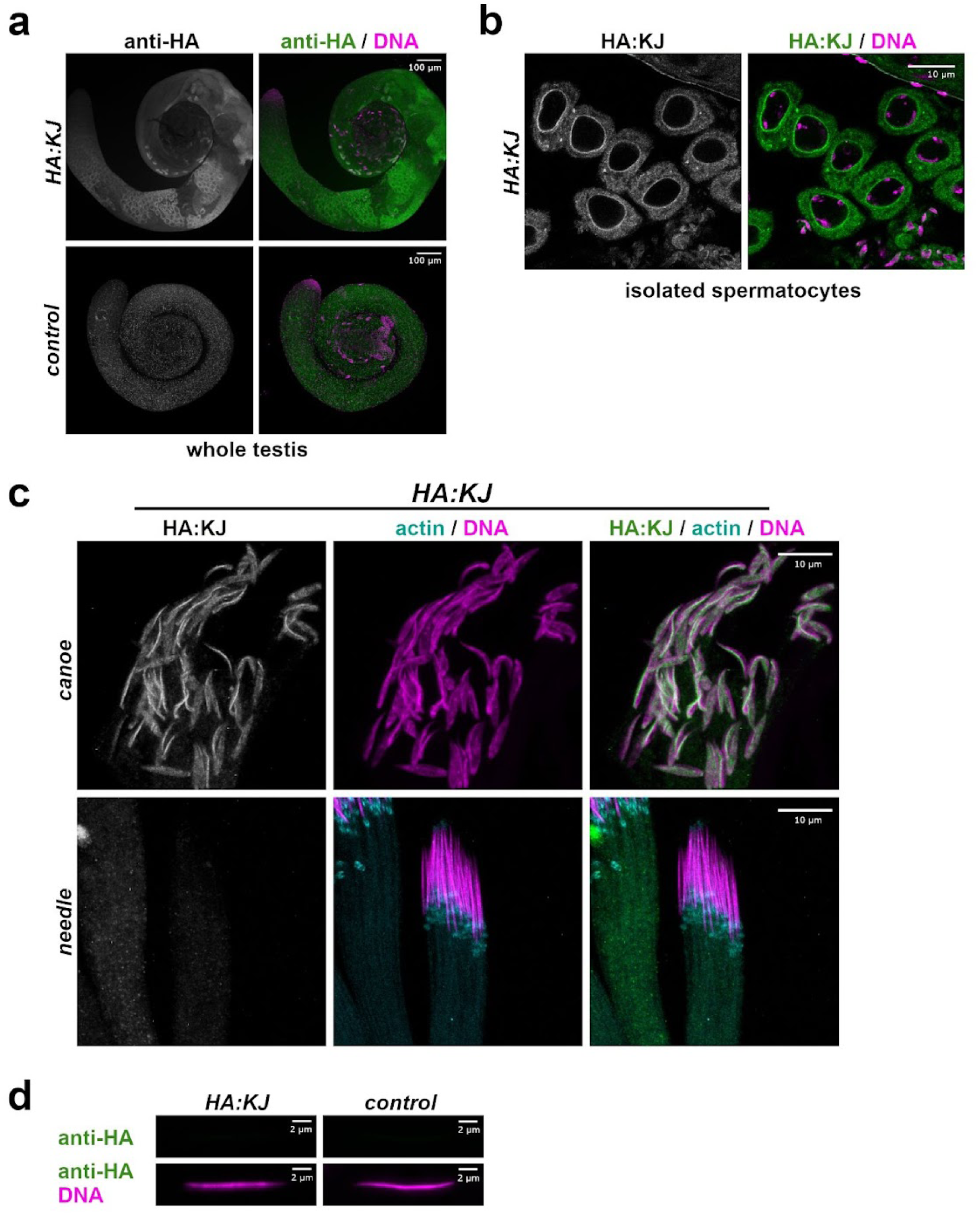
KJ is found around the edge of nuclei in both spermatocytes and spermatids but is undetectable in mature sperm. a) In whole mount testes, HA:KJ (full genotype: Δ*kj*/Δ*k; HA:KJ/+*) is enriched in spermatocytes and condensing spermatid nuclei. A low level of background is present *w*^1118^ control testes stained with anti-HA. b) In isolated spermatocytes, HA:KJ has a diffuse localization throughout the cytoplasm but is enriched at the edge of the nucleus and at large punctate structures of unknown identity. c) In canoe-stage spermatid nuclei, HA:KJ is localized to condensing nuclei, with an enrichment on one side of each nucleus reminiscent of dense bodies. By the needle stage of condensation, marked by the presence of actin-rich investment cones at the base of nuclei, HA:KJ is no longer detectable around nuclei. d) Staining of mature sperm isolated from seminal vesicles shows no detectable HA:KJ around sperm nuclei. Control sperm are from *w*^1118^ males.

### Predicted biochemical properties of KJ protein

The *D. melanogaster kj* gene is located on chromosome 2L (Muller element B), and its single exon is predicted to encode a 126-amino acid protein of predicted molecular weight 15 kDa and a predicted isoelectric point of 8.7. DeepTMHMM (Hallgren et al. 2022) predicts the protein to have one transmembrane domain spanning residues 21-36, with the N-terminus predicted to be outside the membrane and the C-terminus predicted to be inside. AlphaFold3 (Abramson et al. 2024) predicts the protein to have two prominent alpha helices predicted with either very high (pLDDT > 90) or high (70 > pLDDT > 90) confidence: one spanning residues 2-61, and another spanning residues 85-106 (Fig. 5A). Most other regions are predicted to be disordered at a lower confidence level. The DeepLoc 2.1 algorithm (Ødum et al. 2024) predicts that the KJ protein localizes to the endoplasmic reticulum with a 0.92 probability (the prediction probabilities to all other locations were < 0.4).

**Figure 5.**
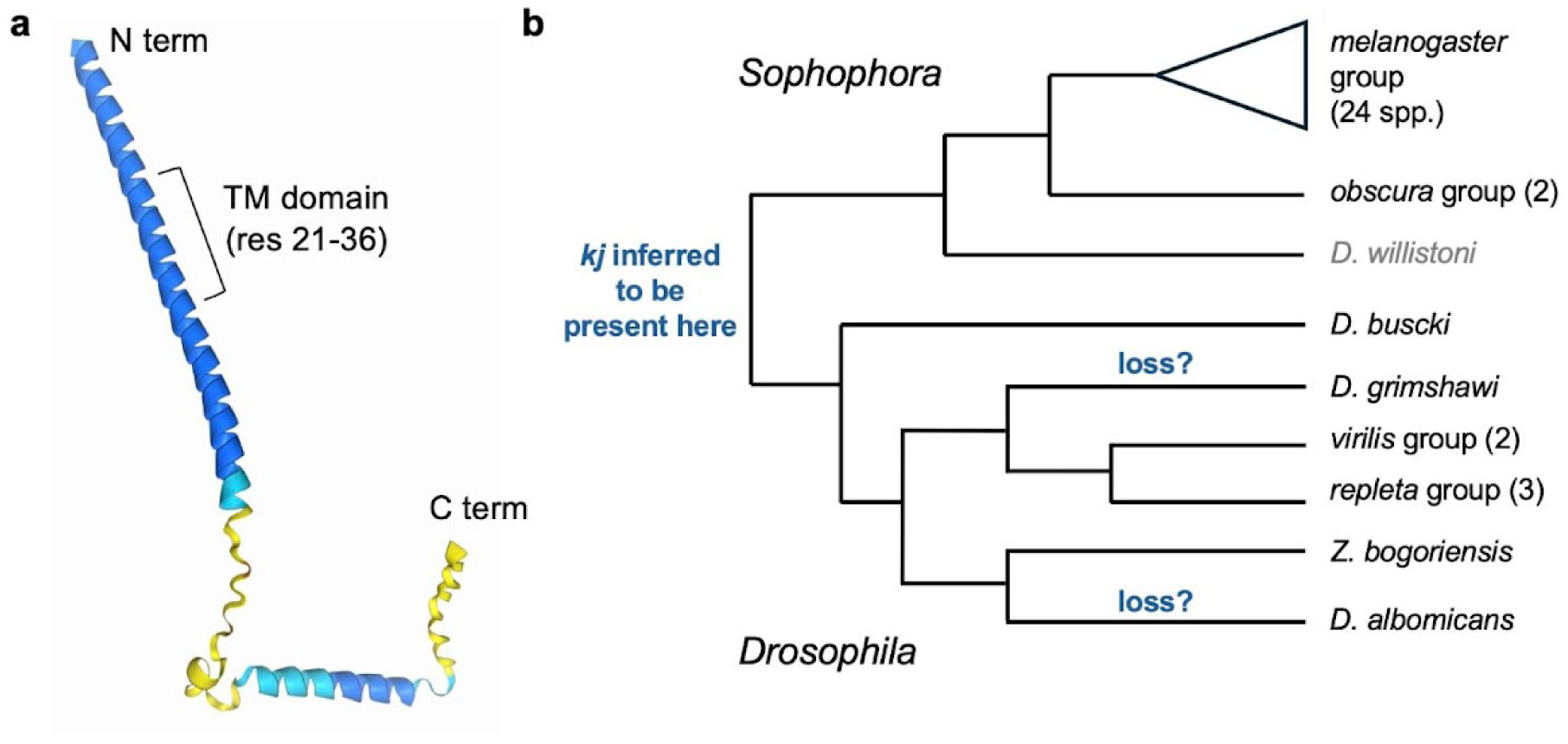
**Predicted KJ protein structure and molecular evolution of *kj* in *Drosophila.*** a) AlphaFold3-predicted structure of the 126-residue *D. melanogaster* KJ protein. The position of the predicted transmembrane (TM) domain is. Color indicates the degree of model confidence (dark blue: very high confidence, pLDDT > 90; light blue: high confidence, 90 > pLDDT > 70; yellow: low confidence, 70 > pLDDT > 50). b) Phylogenetic distribution and potential gene loss events for *kj* in genus *Drosophila*. Orthologs of *kj* were detected in both subgenera, *Sophophora* and *Drosophila*, but not outside of genus *Drosophila*, implying that *kj* was present at the base of the genus. The lack of detectable, intact orthologs in the syntenic regions of the genomes of *D. grimshawi* and *D. albomicans* suggests potential gene loss events in these lineages. Gray text indicates uncertainty about the ortholog identified in *D. willistoni.* For clarity, some species are collapsed into groups; the number of species from the group for which full-length orthologs were detected is shown in parentheses. Branch lengths are not to scale; tree topology shows species relationships inferred by Suvorov et al. (2022).

### Molecular evolution of *kj* in *melanogaster* group species

Because of its lack of identifiable homologs outside of *Drosophila* and lack of identifiable protein domains, the *kj* gene and its encoded protein were characterized as putatively *de novo* evolved in a previous bioinformatic analysis (Heames et al. 2020). Further support for the gene’s *de novo* status came from a comprehensive investigation of *de novo* genes in *D. melanogaster*, which used a whole-genome alignment approach to assess the age of each gene (Peng and Zhao 2024). Both analyses determined that *kj* was restricted to the *melanogaster* group of the *Drosophila* genus (Fig. 5B). Consistent with these results and with expectations for an orphan or *de novo* gene, our BLASTP and iterative PSI-BLAST searches showed no detectable homology to any other protein. PSI-BLAST (and subsequent manual annotation of hits) identified 22 additional full-length orthologs throughout the *melanogaster* group, but not outside of it (Table S1, File S1). We identified partially annotated ortholog fragments in four additional species. Another species, *D. eugracilis*, initially appeared to have a pseudogenized copy of *kj* due to a 1-nucleotide insertion in the ORF, but upon manual inspection we found that this nucleotide was not present in RNA-seq reads that mapped to this location and thus likely represented an error in the reference genome. Based on TimeTree estimates (Kumar et al. 2022), these results would suggest the gene arose ∼25-30 million years ago in the common ancestor of this group. RNA-seq data (Brown et al. 2014; Chen et al. 2014) were available through the Genomics Education Partnership for 16 of the 24 species with putatively functional, full-length orthologs. All 16 of these orthologs are expressed in adult males, and nearly all in a male-specific or heavily male-biased pattern (Table S1). AlphaFold3 modeling of KJ from a diverged, in-group ortholog from *D. ananassae* showed a fairly similar structure to that of *D. melanogaster* KJ, with two prominent alpha helices at similar positions (Fig. 5A and Fig. S6).

Taken together, the expression and structural data suggest *kj* may function in male reproduction across the *melanogaster* group.

Genes that mediate reproduction often evolve at elevated rates (Wilburn and Swanson 2016). We therefore used an alignment of 22 *melanogaster* group orthologs (Table S1; Fig. S7; File S3) to examine the molecular evolution of the *kj* protein-coding sequence and to ask whether any KJ residues had experienced recurrent adaptive evolution. PAML model M0 (Yang et al. 2000) estimated the overall *d*_N_/*d*_S_ ratio across the whole gene as 0.42. When similar whole-gene *d*_N_/*d*_S_ estimates were calculated genome-wide for six representative species of the *melanogaster* group (Drosophila 12 Genomes Consortium et al. 2007; Chang et al. 2023), a value of 0.42 fell into the top 1-2%, suggesting that *kj* evolves more rapidly than most *D. melanogaster* genes.

When we asked whether specific residues of the KJ protein had experienced adaptive evolution, the results were ambiguous. The PAML sites test (Yang et al. 2000) compares the likelihood of a model of molecular evolution (M7) that allows only purifying and neutral evolution to a model (M8) that additionally allows a subset of sites to evolve adaptively with *d*_N_/*d*_S_ > 1. This test found no difference in likelihood between the models (ꭓ^2^ = 0, 2 df, *p* = 1.00) and thus found no evidence of recurrent, adaptive evolution on any KJ residue. An analogous method to detect this type of recurrent selection, the Fixed Effects Likelihood (FEL) analysis in the DataMonkey suite of programs (Kosakovsky Pond and Frost 2005), identified three positions (each with *p* < 0.1) in the alignment as having significant evidence for recurrent, adaptive evolution: positions that aligned to residues 56S and 101R in the *D. melanogaster* protein, as well as residues from other species that aligned to a gap between *D. melanogaster* residues 15A and 16F (Fig. S7). The BUSTED-HM algorithm (Murrell et al. 2015; Wisotsky et al. 2020) found no significant evidence for episodic (as opposed to recurrent) positive selection on specific residues.

Consistent with these results, Peng and Zhao (Peng and Zhao 2024) determined, for a different set of *melanogaster* group species, that most non-synonymous substitutions in KJ were non-adaptive. Thus, we conclude that *kj* evolves rapidly, but with only limited evidence for recurrent adaptive evolution on a few of its sites. In spite of its essential function, the gene’s high overall rate of evolution may instead be due to relaxed constraint (Dapper and Wade 2020) on at least some portions of the protein, as has been observed for a high fraction of fly seminal proteins (Patlar et al. 2021). Inspection of KJ amino acid alignment (Fig. S7) showed that the highest conservation between *melanogaster* group orthologs was found just before and around the prominent alpha helix near the C-terminus and, to a lesser extent, around the predicted transmembrane domain. It is possible that these regions are of heightened functional importance in this group of species.

### Identification of potential *kj* orthologs outside of the *melanogaster* group

*Strong evidence for* kj *orthologs in the* Drosophila *subgenus.* Our previous studies of putative *de novo* genes (Gubala et al. 2017; Rivard et al. 2021) have sometimes identified more distantly related orthologs that were not detectable by BLAST and/or not previously annotated as genes. To investigate the possibility of such orthologs for *kj*, we queried *Drosophila* genomes outside of the *melanogaster* group using TBLASTN with relaxed parameters (e-value threshold < 10, word size = 3). Any hits from these searches were evaluated for their genomic location, their expression pattern based on available RNA-seq data, and whether the inferred potential protein showed homology to *D. melanogaster* KJ. This process identified a potential *kj* ortholog in a *virilis* group species of subgenus *Drosophila*, *D. virilis* (Fig. S8). The initial TBLASTN search identified a 75-nt stretch in this species predicted to encode 25 amino acids with 52% identity (72% similarity) to a region of *D. melanogaster* KJ, producing an e-value of 8.3. This hit’s position in the *D. virilis* genome is syntenic to the position of *kj* in *D. melanogaster* because it is flanked by three of the same genes that surround *kj* in *D. melanogaster* (orthologs of *CG6614* and *CG4983* upstream, and the ortholog of *Vha100-5* downstream). The region identified by TBLASTN exists within a potential open reading frame (ORF) that could encode 171 amino acids. The genomic region encoding this ORF showed signals of expression in RNA-seq data from both sexes of adult *D. virilis*. The maximum read depth was 43-fold higher in males, consistent with a gene that functions in male reproduction. A pairwise BLASTP comparison of the full *D. virilis* ORF to *D. melanogaster* KJ produced a significant e-value of 10^-7^, and DeepTMHMM predicted a single transmembrane domain with the same orientation with respect to the membrane (N terminus outside, C terminus inside) as *D. melanogaster* KJ. A small, duplicated amino acid motif in the C terminus of the putative *D. virilis* ortholog contributes to this ortholog’s longer length (Fig. S8).

The presence of a likely *kj* ortholog in the *Drosophila* subgenus implied that the origin of the *kj* gene could be earlier than the previously estimated 25-30 million years ago. To determine the phylogenetic distribution of *kj* across the genus, we used a combination of BLASTP, TBLASTN and synteny to search for additional orthologs in a variety of species and groups (Fig. 5B).

These methods identified proteins of similar length and the same DeepTMHMM-predicted topology in another *virilis* group species, *D. novamexicana*; three members of the *repleta* group species (*D. hydei, D. mojavensis, D. arizonae*); and, additional species *D. busckii* and *Zaprionus bogoriensis* (File S2); *Zaprionus* is a genus within the paraphyletic *Drosophila* genus (see Fig. 5B.) The syntenic region of *repleta* group member *D. navojoa* contained a much shorter ORF (60 a.a.) with male-specific expression and sequence identity to these orthologs, but the predicted protein did not contain a transmembrane domain, so this region may represent a pseudogene or a gene with altered function. Table S2 lists the genomic locations and biochemical properties of the likely orthologs outside of the *melanogaster* group. AlphaFold3 modeling of representative orthologs from each lineage produced predicted structures that were fairly similar to those of representative *melanogaster* group orthologs, with a long alpha helix predicted with high confidence toward the N terminus of each ortholog and one or two shorter alpha helices in the C-terminal half of the protein (Fig. S6).

#### Somewhat strong evidence for kj orthologs in the obscura group

Since we detected *kj* orthologs in both the *Sophophora* and *Drosophila* subgenera, we wondered whether *kj* was present in the *obscura* group, a part of the *Sophophora* subgenus distinct from the *melanogaster* group (Fig. 5B). Using *D. pseudoobscura* and *D. subobscura* as representative species, we identified in their syntenic regions ORFs supported as male-expressed by RNA-seq data that could encode proteins of similar length to *D. melanogaster* KJ (Table S2). These ORFs were predicted by DeepTMHMM to have a single transmembrane domain in the same approximate position as the KJ orthologs described above, though the predicted topology (N terminus inside the membrane, C terminus outside) was inverted. The predicted proteins showed significant identity to each other across their full lengths. Pairwise BLASTP homology to the above-detected KJ orthologs was marginal. The *D. pseudoobscura* ORF, for example, matched three orthologs from the *melanogaster* (*D. erecta*, *D. setifemur*) and *repleta* groups (*D. arizonae*) with 0.01 < e < 0.05, and sixteen other orthologs with e < 5. Most of these matches corresponded to the predicted transmembrane domain. The data were similar for *D. subobscura*: its predicted ORF produced BLAST hits to nine other KJ orthologs with e-values ranging from 0.003 to 0.53, with most regions of sequence identity falling in the predicted transmembrane domain. AlphaFold3 modeling of the *D. pseudoobscura* ORF showed a broadly similar structure to other KJ orthologs (Fig. S6), increasing confidence that these ORFs could represent true *kj* orthologs.

#### Levels of amino conservation in non-melanogaster group orthologs

We aligned the above-described KJ orthologs to examine levels of amino acid conservation. Two general regions of heightened conservation were apparent (Fig. S9), both similar in position to the two more highly conserved regions of the *melanogaster* group orthologs (Fig. S7). One region was toward the C-terminus and partially overlapped with a predicted alpha helix in the *D. virilis* ortholog. The other surrounded the predicted transmembrane domain toward the N-terminus.

Overall levels of conservation were somewhat lower for these orthologs, as expected given the wider phylogenetic range represented by the included species (Fig. 5B).

*Marginal evidence for a* kj *ortholog in* D. willistoni. We identified a potential *kj* ortholog in *D. willistoni*, a *Sophophora* subgenus species that is an outgroup to both the *melanogaster* and *obscura* groups, by examining regions with male gonad RNA-seq expression data within the syntenic region (Fig. S10). One such region showed the potential to encode a protein of 138 amino acids, with one predicted transmembrane domain (though only a single residue, the first methionine of the polypeptide, is predicted to be outside the membrane). The full-length ORF had marginal BLASTP similarity to potential KJ orthologs from the *obscura* group (e-values between 0.5 and 1). Its predicted protein structure, however, did not have the same confidently predicted alpha helices as the other orthologs (Fig. S7), and the amino acid sequence did not align well with the other orthologs. Thus, *D. willistoni* may have a *kj* ortholog, but the evidence is ambiguous.

*Inability to detect* kj *in* D. grimshawi *and* D. albomicans. Two remaining *Drosophila* subgenus species for which good RNA-seq and genome browser data were available were *D. grimshawi* and *D. albomicans*. Both of these species are nested within the *Drosophila* subgenus.

TBLASTN searches of the orthologs above against the whole genomes of either species did not produce any meaningful hits, so we focused on the syntenic region. For *D. grimshawi* (Fig. S11), the *D. virilis* ortholog produced a reasonably strong TBLASTN hit within the syntenic region (e < 10^-6^ across a 65-residue region of homology toward the C-terminus of the protein). However, this region had stop codons immediately upstream of it in all three reading frames, and RNA-seq coverage was spotty and at a much lower level than we observed for better-supported orthologs. Thus, we find no evidence of a functional *kj* in the *D. grimshawi* syntenic region; instead, the evidence may be consistent with a somewhat recent pseudogenization event.

For *D. albomicans*, we identified three regions with male-specific/biased expression in the syntenic region (Fig. S11). None were predicted to encode an ORF of >65 amino acids, and none of the potential ORFs had predicted transmembrane domains. When we used TBLASTN to query the entire syntenic region (150,000 bp) for regions of potential homology to any *Drosophila* subgenus KJ ortholog (Table S2), we found no promising hits Thus, we conclude there is no detectable *kj* ortholog in the *D. albomicans* syntenic region.

#### Conclusions about kj age and phylogenetic distribution

Collectively, these phylogenetic data suggest that *kj* was likely present at the base of the *Drosophila* genus, estimated by TimeTree to be ∼43 million years ago (Kumar et al. 2022). Whether the gene originated in the common ancestor of *Drosophila* or more anciently is unclear. Using similar methods to those described for *D. albomicans* and *D. grimshawi*, we investigated the syntenic region of outgroup species *Scaptodrosophila lebanonensis*. This search yielded no obvious *kj* ortholog, but detecting orthologs of short, fast-evolving *Drosophila* genes outside of the genus is challenging (Weisman et al. 2020). Regardless of the exact timing of *kj*’s origin, it appears to be more ancient than previously estimated (Heames et al. 2020; Peng and Zhao 2024). To be prudent, we describe *kj* as an orphan gene (Tautz and Domazet-Lošo 2011), though we note that none of the evidence above is inconsistent with a *de novo* origin for this gene. We use the more cautious “orphan” terminology, however, to account for other possibilities consistent with our inability to detect orthologs outside of *Drosophila*, such as rapid sequence divergence, movement of the gene to a different genomic location, an origin via gene fusion, gene truncation, or horizontal gene transfer, or an apparent absence in outgroup species due to incomplete genome assemblies.

Within genus *Drosophila*, *kj* has been fairly well conserved, but our inability to detect the gene in two lineages (*D. grimshawi* and *D. albomicans*) in which the syntenic region remains intact suggest the possibility of lineage-specific gene loss events (Fig. 5). Additional potential losses, or major changes in protein structure/function, are possible in *D. willistoni* and *D. navojoa*.

Within the *melanogaster* group, however, the gene and its male-specific expression pattern are well conserved.

## Discussion

We have identified a *D. melanogaster* gene, *katherine johnson* (*kj*), whose action is required for efficient sperm entry into eggs. Interestingly, *kj* is an orphan gene that was likely present at the origin of the *Drosophila* genus but has evolved rapidly since then. These findings hold promise for unraveling the still mysterious molecular events surrounding *Drosophila* fertilization and reinforce the idea that lineage-specific genes can evolve essential roles in broadly conserved biological processes.

### Potential functions for KJ in spermatogenesis

Relatively little is known about the molecules required for sperm-egg interactions in *Drosophila* (Loppin et al. 2015). Mutations in several genes result in normal sperm production and transfer, but low hatchability, as we observe for *kj*. However, the cellular causes of their fertility defects are distinct. Mutants in genes like *wasted* and *Nep4* cause abnormal sperm storage or release, resulting in lower rates of fertilization (Ohsako and Yamamoto 2011; Ohsako et al. 2021), but sperm from *kj* nulls appear to be stored and released normally (Table 1). Paternal effect mutants cause abnormalities in processes such as sperm membrane dissolution (Fitch and Wakimoto 1998; Wilson et al. 2006) or paternal chromatin unpacking or reorganization (Loppin, Lepetit, et al. 2005; Dubruille et al. 2023), but sperm from these mutant males are proficient at egg entry, unlike sperm from *kj* nulls (Fig. 3). Thus, *kj* is the only extant and molecularly characterized *Drosophila* gene that distinctly affects sperm entry into eggs.

One other gene, *casanova* (*csn*), had been reported to have a mutant phenotype similar to what we find for *kj*: *csn* mutant males produce and transfer motile, morphologically normal sperm that are stored properly, but are unable to fertilize eggs (Perotti et al. 2001). Unfortunately, *csn* mutants are no longer available, and the molecular nature of the gene is unknown. It is clear that *csn* is distinct from *kj*, since they map to different chromosomal positions (*kj* is at cytological region 34F4 on chromosome arm 2L; *csn* was mapped to cytological region 95E8-F7 on chromosome arm 3R). It has been proposed that sperm interact with and/or cleave β-N-acetylglucosamine and ɑ-mannose sugars that are present on the egg at the site of sperm entry but are no longer detected after fertilization (Loppin et al. 2015). Sperm plasma membranes have been reported to contain glycosidic enzymes that cleave these sugars (Cattaneo et al. 2002; Intra et al. 2006), and sperm β-N-acetylglucosaminidase activity is reduced in *csn* mutants (Perotti et al. 2001). Our data suggest KJ is not involved in such carbohydrate interactions between egg and sperm, as it is not detected by immunofluorescence on mature sperm from seminal vesicles (Fig. 4D). KJ was also not detected in the mature sperm proteome as determined by mass spectrometry (Garlovsky et al. 2022). While we recognize the limitations to negative results with both of these detection methods, the lack of any sequence similarity of KJ to any glycolytic enzyme supports our view that KJ is unlikely to participate directly in sperm-egg carbohydrate interactions.

The localization pattern of HA:KJ in the testes (Fig. 4) allows us to hypothesize different potential roles for the KJ protein in spermatogenesis. In spermatocytes, KJ is enriched around the entire edge of the nucleus, with fainter staining visible throughout the cytoplasm. In spermatids, however, KJ localizes along one side of the elongating nuclei. These patterns could be consistent with three possible functions for KJ. First, as predicted by DeepLoc, KJ may localize to all or a portion of the ER, where it could be embedded in the membrane via its predicted transmembrane domain. ER localization is consistent with the observed pattern of HA:KJ in spermatocytes and spermatids (Fig. 4). In spermatocytes, the ER is continuous with the outer nuclear membrane and extends into the cytoplasm (Lindsley and Tokuyasu 1980; Dorogova et al. 2009), matching the HA:KJ localization pattern (Fig. 4B). As spermatid nuclei begin to condense after meiosis, a portion of the ER remains associated with the one edge of the nuclear membrane (Lindsley and Tokuyasu 1980), consistent with our observation of HA:KJ along a single edge of the nucleus at this stage (Fig. 4C). Later, during individualization, the ER (and other organelles) are stripped from the spermatids by individualization complexes and discarded in waste bags at the apical end of the spermatid cyst discarded (Dorogova et al. 2009). The removal of the ER during the final stages of spermiogenesis is consistent with the absence of detectable HA:KJ in mature sperm. Under this scenario, the inability of sperm from *kj* null males to enter eggs could potentially be caused by the loss of KJ protein from a key organelle for protein folding, processing and transport, which could in turn result in defects in the production and/or transport of components of the mature sperm proteome that are necessary for efficient egg entry.

A second possibility, consistent with HA:KJ’s localization in spermatids (Fig. 4C), is that KJ could be a component of the dense body. This structure, analogous to the mammalian manchette (Lehti and Sironen 2016), forms through close physical interactions between nuclear membrane proteins, microtubules, and actin-based structures and that maintains contact between the condensing spermatid nuclei and microtubules that help form the elongating sperm tail (Fabian and Brill 2012). Unlike *kj* mutants, however, mutations in genes that alter dense body formation cause defects in nuclear shaping at late stages of spermatogenesis, blocking mature sperm production and resulting in complete sterility (Kracklauer et al. 2010; Augière et al. 2019; Li et al. 2023) For example, a recently characterized protein, Mst27D, appears to function in dense body formation and nuclear shaping, as it physically links nuclear pore complex proteins with microtubules (Li et al. 2023). As loss of *kj* has no effect on nuclear shaping, *kj* most likely is not required for dense body formation and therefore is likely to act independently of *Mst27D*. Instead, if KJ localizes to the dense body, it might implicate roles for the structure beyond nuclear and sperm head shaping, possibly in sperm head organization and protein trafficking.

Third, KJ’s localization around the entire nucleus in spermatocytes and along one edge of the nucleus in spermatids could be consistent with the protein functioning in or adjacent to the nuclear membrane. Although we do not see gross changes in sperm nucleus/head shape in the absence of *kj*, its loss might cause subtle abnormalities in these regions that make it more difficult for sperm to enter the micropyle, the size of which coevolves with the diameter of insect sperm (Soulsbury and Iossa 2024). Alternatively, it is possible that *kj* mediates (through either nuclear membrane or ER localization) some other aspect of sperm head organization, such as ensuring correct localization of other proteins, or acts in another process required to prepare sperm for efficient egg entry or to release sperm from storage in a way that facilitates their interaction with the egg.

### KJ molecular evolution

Consistent with the analysis of Peng and Zhao (2024), we found that the *kj* gene is well conserved in the *melanogaster* group. We also observed that these orthologs show strongly male-biased expression. This pattern is consistent with the hypothesis that *kj* may play an important role in male reproduction across species in this clade and, thus, that it might have already evolved its essential function in the common ancestor of this group. However, the *kj* protein-coding sequence has evolved considerably faster than most genes do in this group of species, with limited evidence of recurrent adaptation. This pattern could indicate that only some regions of the KJ protein are important for its essential function (while others evolve under relaxed constraint), consistent with the differing levels of conservation that we observed in the aligned orthologs (Figs. S7 and S9), and/or that the protein’s essential function arose in a more recent ancestor of *D. melanogaster*. *kj* was initially identified as a putative *de novo* gene because of the lack of detectable orthologs outside of *Drosophila* and the lack of identifiable protein domains (Heames et al. 2020). A sophisticated analysis using whole-genome alignments similarly concluded that *kj* was unique to *melanogaster* group species (Peng and Zhao 2024). Since our approach to ortholog detection was tailored to the *kj* gene, we were able to use features specific to *kj* (such as synteny, expression pattern and predicted protein features) and a relaxed threshold for initial BLAST searches to identify potential *kj* orthologs beyond the *melanogaster* group. This gene-specific approach would not have been feasible for the previous genome-scale studies. Our results highlight the utility of considering gene-specific parameters when searching for orthologs of putative *de novo* genes and suggest that caution may be warranted when a gene’s *de novo* status is supported only by high-throughput bioinformatic analysis.

While we found *kj* orthologs broadly across the *Drosophila* genus, we also found several lineages in which the gene was either undetectable, truncated, or predicted to encode a protein with a drastically different predicted structure from *D. melanogaster* KJ. Thus, while *kj* was likely present in the *Drosophila* common ancestor, it might have been dispensable in some lineages. The phylogenetic distribution of *kj* is similar to the distributions of two other orphan (previously termed “putative *de novo*”) genes with essential male reproductive functions in *D. melanogaster*, *saturn* and *atlas* (Gubala et al. 2017; Rivard et al. 2021), which are also well conserved in the *melanogaster* group and detectable in only some outgroup species. Our general hypothesis for this pattern is that these genes could have had slight, positive effects on fertility in the most ancient ancestors of the *Drosophila* genus before evolving more essential, non-redundant roles in the lineage leading to the *melanogaster* group. It is also possible that larger-scale changes to the process of spermatogenesis in specific lineages could have rendered once-beneficial genes superfluous. Indeed, several instances of major, lineage-specific changes in spermatogenesis are known, such as the evolution of three types of sperm, only one of which is fertilization competent, in *D. pseudoobscura* and related species (Alpern et al. 2019), and the evolution of new sex chromosomes, which can affect processes such as the regulation of sex-linked genes in germline cells (Wei et al. 2024) and sex chromosome meiotic drive (Chang et al. 2023).

While we identified likely *kj* orthologs across *Drosophila* species, neither BLAST, PSI-BLAST nor HMMER (hmmer.org) could detect homologs outside the genus, and we could not identify a homolog in the syntenic region of *S. lebanonensis*. Thus, the phylogenetic distribution of *kj* currently appears to be restricted to genus *Drosophila*. Because there is no evidence that *kj* arose through duplication, we consider it an orphan gene (Tautz and Domazet-Lošo 2011). It is possible that more sensitive sequence- or structure-based methods will at some point identify a *kj* ortholog outside of *Drosophila*. Even if such an ortholog is detected, however, a gene that is required for efficient fertilization and that has evolved within *Drosophila* to the point that it is currently unrecognizable in outgroup species would remain of considerable functional and evolutionary interest. This study provides another demonstration of the important reproductive roles that lineage-specific genes can evolve, in this case in the crucial process of sperm entry into eggs in *D. melanogaster*. As genome editing becomes easier to perform in non-model species (Bubnell et al. 2022), it should also be possible to test whether and how *kj* is required for male fertility in other *Drosophila* species.

## Data Availability Statement

Fly strains are available on request. Files S1 and S2 contain the inferred protein sequences of predicted KJ orthologs. File S3 contains the DNA sequence alignment used in the molecular evolutionary analyses. File S4 contains the phylogenetic tree used for PAML analysis. File S5 contains the raw data underlying this study’s graphs and statistical analyses. Genome browsers and RNA-seq data for *Drosophila* species were accessed through the Genomics Education Partnership (thegep.org). Other supporting information is provided in either the supplemental figures and tables or in the Reagents Table.

## Supporting information

Reagents Table

## Acknowledgements

We thank Biounce Casildo Ramirez, Roselyn Pantoja-Ramirez, Rob Bellin and Karen Ober and for technical assistance; Wilson Leung for assistance for checking the *D. eugracilis* RNA-seq reads; Marcus Kilwein for assistance with embryo staging; and Barbara Wakimoto, Emily Rivard, Lars Eicholt, Harmit Malik and members of the Findlay and Wolfner Labs for feedback on the project. The reviewers and editor provided helpful feedback on the manuscript. This work was supported by NSF grants 1652013 and 2212972 to GDF. JMT was supported by a Mann Fellowship (to JMT) and Payne Memorial Funds (to MFW). MFW also acknowledges support from NIH grant R37-HD038921. KLM was supported by a College of the Holy Cross Weiss summer research fellowship.

## List of Supplemental Material

**Figure S1: RNAi reagents and assessment of knockdown.** A) Oligos to create a hairpin siRNA to be cloned into pValium20, which created a siRNA targeting the *CG43167* gene. B) RT-PCR of cDNA from whole males that were either knocked down for *kj* (*Bam*-GAL4, UAS-*Dicer2* > UAS:*CG43167* RNAi) or control (*Bam*-GAL4, UAS-*Dicer2*) shows that *kj* transcript levels are essentially undetectable in knockdown males. C) RT-PCR of *RpL32* is shown as a positive control for successful cDNA synthesis. D) RT-PCR primers for detecting *kj* expression.

**Figure S2: CRISPR reagents and confirmation of *kj* deletion allele.** A) Chromosomal coordinates of guide RNA (gRNA) target sites flanking *CG43167*. Coordinates of the region that was ultimately deleted in the Δ*kj* allele are shown. The deletion was initially detected by PCR screening using the primers indicated. B) UCSC genome browser graphic showing the BLAT result when the primers were used to amplify and sequence genomic DNA from Δ*kj* homozygous males. The entire *kj*/*CG43167* gene is deleted.

**Figure S3: Gross testis morphology of *kj* null males appears normal under phase contrast microscopy.** A) Testis and seminal vesicle from a *kj*+/*kj*+ male. B) Testis and seminal vesicle from a Δ*kj*/Δ*kj* male.

**Figure S4: Epifluorescence images used to quantify presence of Dj-GFP in embryos.** Representative epifluorescence images used to quantify presence of Dj-GFP in Fig. 4C. Cyan arrows indicate sperm marked by dj-GFP. Scale bar = 100μm.

**Figure S5: HA:KJ is not detected in mature sperm nuclei that underwent decondensation treatment.** Sperm isolated from seminal vesicles were first pre-treated with 1X PBS supplemented with 1% Triton X-100 for 30 minutes. They were then treated with a decondensation buffer. Staining with anti-HA did not detect HA:KJ around the nucleus of either pre-treated (A) or decondensed (B) sperm. While this experiment lacked a positive control, the intensity and shape of the DNA signal changed in the decondensed sperm, suggesting at least some degree of effective decondensation.

**Figure S6: AlphaFold3 protein structure predictions of KJ orthologs.** The top AlphaFold3-predicted structure for *Drosophila melanogaster* KJ and the identified orthologs from 1. *D. ananassae*, *D. pseudoobscura*, *D. willistoni*, *D. virilis*, *D. mojavensis*, *Zaprionus bogoriensis* and *D. buscki*. In each case, the structure was rotated so as to position the N terminus at the top of the image. Colors indicate pIDDT level, a measure of the degree of confidence in the position of each residue.

**Figure S7: Amino acid alignment of KJ orthologs from the *melanogaster* group.** KJ protein sequences from *melanogaster* group orthologs were aligned using ClustalOmega and visualized in Jalview. The positions of the predicted alpha helices (black boxes) and transmembrane domain (blue line) from the *D. melanogaster* ortholog are shown above the alignment. Aligned positions identified by FEL as being under positive selection are indicated by asterisks at the bottom of the alignment.

**Figure S8: Evidence for a *kj* ortholog in *Drosophila virilis.*** A) In *D. melanogaster*, the *kj* gene (CG43167) is flanked by three genes that are well conserved in other *Drosophila* species: *CG6614, CG4983* and *Vha100-5*. B) The syntenic region of the *D. virilis* genome contains orthologs of these same three genes. The orange box indicates the stretch within the syntenic region that had faint TBLASTN homology to *D. melanogaster* KJ. C) TBLASTN hit details and RNA-seq evidence of male-biased expression at the putative orthologous region of *D. virilis*. D) The full-length ORF surrounding the *D. virilis* TBLASTN hit (original peptide matching *D. melanogaster* KJ is in orange; two potential start codons are in green). The full-length *D. virilis* ORF has significant identity to *D. melanogaster* KJ in a pairwise BLASTP comparison. This comparison shows a small region of potential duplication toward the end of the *D. virilis* ORF. The pairwise BLASTP gave a significant e-value while the genome-wide TBLASTN did not because the latter considered a much larger set of possible subject sequences. E) DeepTMHMM predictions of transmembrane domains and protein topology for KJ in *D. melanogaster* (left) and the putative *D. virilis* ortholog (right).

**Figure S9: Amino acid alignment of KJ orthologs from outside of the *melanogaster* group.** KJ protein sequences from orthologs identified outside of the *melanogaster* group were aligned using ClustalOmega and visualized in Jalview. The positions of the predicted alpha helices (black boxes) and transmembrane domain (blue line) from the *D. virilis* ortholog are shown above the alignment. The putative ortholog from *D. willistoni* aligned poorly and was thus excluded.

**Figure S10: Evidence for a possible *kj* ortholog in *Drosophila willistoni.*** A) Unannotated regions of the *kj* syntenic region that showed RNA-seq expression in adult males were evaluated to determine whether they could encode a *kj* ortholg. Most expressed regions were ruled out for reasons listed at bottom left. Region B, however, encoded an ORF with a predicted transmembrane domain that showed marginal BLASTP similarity to *kj* orthologs from the *obscura* group. B-C) Details of RNA-seq expression, transmembrane prediction and BLASTP results of the possible ortholog.

**Figure S11: Evidence for *kj* gene absence in *Drosophila albomicans* and *Drosophila grimshawi***. A) The *kj* syntenic region of *D. albomicans* contains three unannotated regions (A-C) with RNA-seq evidence of male expression. None of these regions is likely to encode a *kj*-like ORF for reasons listed at the bottom left. B) One stretch of the *D. albomicans* syntenic region had very faint TBLASTN homology to *D. virilis* KJ, but the region has no extended ORF and no evidence for adult expression. C) In *D. grimshawi*, TLBASTN identified one stretch of the *kj* syntenic region that has homology to *D. virilis* KJ. There is very low, spotty coverage of this region from adult male RNA-seq data, but no extended ORFs that would include the TBLASTN hit. These data may be consistent with *kj* pseudogenization in *D. grimshawi*.

Table S1: Coordinates, protein lengths, predicted protein topologies, and expression data of putative *kj* orthologs in *melanogaster* group species.

Table S2: Coordinates, protein lengths, predicted protein topologies, and expression data of putative *kj* orthologs outside of the *melanogaster* group.

File S1: Protein sequences of *kj* orthologs from *melanogaster* group *Drosophila* species.

File S2: Protein sequences from putative *kj* orthologs from non-*melanogaster* group *Drosophila* species.

File S3: Alignment of protein-coding DNA sequences of *kj* orthologs from *melanogaster* group *Drosophila* species.

File S4: Phylogenetic tree of protein-coding DNA sequences of *kj* orthologs from *melanogaster* group *Drosophila* species.

File S5: Data underlying the statistics and graphs in Figures 1-3 and Table 1. Reagents Table

## References

1. Abramson J, Adler J, Dunger J, Evans R, Green T, Pritzel A, Ronneberger O, Willmore L, Ballard AJ, Bambrick J, et al. 2024. Accurate structure prediction of biomolecular interactions with AlphaFold 3. Nature. 630(8016):493–500.

2. Alpern JHM, Asselin MM, Moehring AJ. 2019. Identification of a novel sperm class and its role in fertilization in Drosophila. J Evol Biol. 32(3):259–266.

3. Augière C, Lapart J-A, Duteyrat J-L, Cortier E, Maire C, Thomas J, Durand B. 2019. salto/CG13164 is required for sperm head morphogenesis in Drosophila. Mol Biol Cell. 30(5):636–645.

4. Begun DJ, Lindfors HA, Kern AD, Jones CD. 2007. Evidence for de novo evolution of testis-expressed genes in the Drosophila yakuba/Drosophila erecta clade. Genetics. 176(2):1131–1137.

5. Bloch Qazi MC, Heifetz Y, Wolfner MF. 2003. The developments between gametogenesis and fertilization: ovulation and female sperm storage in Drosophila melanogaster. Dev Biol. 256(2):195–211.

6. Brown JB, Boley N, Eisman R, May GE, Stoiber MH, Duff MO, Booth BW, Wen J, Park S, Suzuki AM, et al. 2014. Diversity and dynamics of the Drosophila transcriptome. Nature. 512(7515):393–399.

7. Bubnell JE, Ulbing CKS, Fernandez Begne P, Aquadro CF. 2022. Functional Divergence of the bag-of-marbles Gene in the Drosophila melanogaster Species Group. Mol Biol Evol. 39(7):msac137.

8. Carvunis A-R, Rolland T, Wapinski I, Calderwood MA, Yildirim MA, Simonis N, Charloteaux B, Hidalgo CA, Barbette J, Santhanam B, et al. 2012. Proto-genes and de novo gene birth. Nature. 487(7407):370–374.

9. Cattaneo F, Ogiso M, Hoshi M, Perotti ME, Pasini ME. 2002. Purification and characterization of the plasma membrane glycosidases of Drosophila melanogaster spermatozoa. Insect Biochem Mol Biol. 32(8):929–941.

10. Chang C-H, Mejia Natividad I, Malik HS. 2023. Expansion and loss of sperm nuclear basic protein genes in Drosophila correspond with genetic conflicts between sex chromosomes. Elife. 12:e85249.

11. Chen Z-X, Sturgill D, Qu J, Jiang H, Park S, Boley N, Suzuki AM, Fletcher AR, Plachetzki DC, FitzGerald PC, et al. 2014. Comparative validation of the D. melanogaster modENCODE transcriptome annotation. Genome Res. 24(7):1209–1223.

12. Dapper AL, Wade MJ. 2020. Relaxed Selection and the Rapid Evolution of Reproductive Genes. Trends Genet. 36(9):640–649.

13. Deneke VE, Pauli A. 2021. The Fertilization Enigma: How Sperm and Egg Fuse. Annu Rev Cell Dev Biol. 37:391–414.

14. Ding Y, Zhao L, Yang S, Jiang Y, Chen Y, Zhao R, Zhang Y, Zhang G, Dong Y, Yu H, et al. 2010. A young Drosophila duplicate gene plays essential roles in spermatogenesis by regulating several Y-linked male fertility genes. PLoS Genet. 6(12):e1001255.

15. Dorogova NV, Nerusheva OO, Omelyanchuk LV. 2009. Structural organization and dynamics of the endoplasmic reticulum during spermatogenesis of Drosophila melanogaster: Studies using PDI-GFP chimera protein. Biochemistry (Moscow) Supplement Series A: Membrane and Cell Biology. 3(1):55–61.

16. Drosophila 12 Genomes Consortium, Clark AG, Eisen MB, Smith DR, Bergman CM, Oliver B, Markow TA, Kaufman TC, Kellis M, Gelbart W, et al. 2007. Evolution of genes and genomes on the Drosophila phylogeny. Nature. 450(7167):203–218.

17. Dubruille R, Herbette M, Revel M, Horard B, Chang C-H, Loppin B. 2023. Histone removal in sperm protects paternal chromosomes from premature division at fertilization. Science. 382(6671):725–731.

18. Dubruille R, Orsi GA, Delabaere L, Cortier E, Couble P, Marais GAB, Loppin B. 2010. Specialization of a Drosophila Capping Protein Essential for the Protection of Sperm Telomeres. Current Biology. 20(23):2090–2099.

19. Elofsson A, Han L, Bianchi E, Wright GJ, Jovine L. 2024. Deep learning insights into the architecture of the mammalian egg-sperm fusion synapse. Elife. 13:RP93131.

20. Eren-Ghiani Z, Rathke C, Theofel I, Renkawitz-Pohl R. 2015. Prtl99C Acts Together with Protamines and Safeguards Male Fertility in Drosophila. Cell Rep. 13(11):2327–2335.

21. Fabian L, Brill JA. 2012. Drosophila spermiogenesis: Big things come from little packages. Spermatogenesis. 2(3):197–212.

22. Findlay GD, MacCoss MJ, Swanson WJ. 2009. Proteomic discovery of previously unannotated, rapidly evolving seminal fluid genes in Drosophila. Genome Res. 19(5):886–896.

23. Findlay GD, Sitnik JL, Wang W, Aquadro CF, Clark NL, Wolfner MF. 2014. Evolutionary rate covariation identifies new members of a protein network required for Drosophila melanogaster female post-mating responses. PLoS Genet. 10(1):e1004108.

24. Fitch KR, Wakimoto BT. 1998. The paternal effect gene ms(3)sneaky is required for sperm activation and the initiation of embryogenesis in Drosophila melanogaster. Dev Biol. 197(2):270–282.

25. Foe VE, Odell GM, Edgar BA. 1993. Mitosis and morphogenesis in the Drosophila embryo: point and counterpoint. In: Bate M, Martinez Arias A, editors. The Development of Drosophila melanogaster. Vol. 1. Cold Spring Harbor Laboratory Press. p. 149–300.

26. Garlovsky MD, Sandler JA, Karr TL. 2022. Functional Diversity and Evolution of the Drosophila Sperm Proteome. Mol Cell Proteomics. 21(10):100281.

27. Ge DT, Tipping C, Brodsky MH, Zamore PD. 2016. Rapid Screening for CRISPR-Directed Editing of the Drosophila Genome Using white Coconversion. G3 . 6(10):3197–3206.

28. Gibson DG, Young L, Chuang R-Y, Venter JC, Hutchison CA 3rd, Smith HO. 2009. Enzymatic assembly of DNA molecules up to several hundred kilobases. Nat Methods. 6(5):343–345.

29. Gubala A, Schmitz JF, Kearns MJ, Vinh T, Bornberg-Bauer E, Wolfner M, Findlay GD. 2017. The Goddard and Saturn genes are essential for Drosophila male fertility and may have arisen DE Novo. Mol Biol Evol. 34:1066–1082.

30. Hallgren J, Tsirigos KD, Pedersen MD, Armenteros JJA, Marcatili P, Nielsen H, Krogh A, Winther O. 2022. DeepTMHMM predicts alpha and beta transmembrane proteins using deep neural networks. bioRxiv.:2022.04.08.487609. doi:10.1101/2022.04.08.487609. [accessed 2024 Jul 25]. https://www.biorxiv.org/content/biorxiv/early/2022/04/10/2022.04.08.487609.

31. Heames B, Schmitz J, Bornberg-Bauer E. 2020. A Continuum of Evolving De Novo Genes Drives Protein-Coding Novelty in Drosophila. J Mol Evol. 88(4):382–398.

32. Horne-Badovinac S. 2020. The Drosophila micropyle as a system to study how epithelia build complex extracellular structures. Philos Trans R Soc Lond B Biol Sci. 375(1809):20190561.

33. Horner VL, Wolfner MF. 2008. Transitioning from egg to embryo: triggers and mechanisms of egg activation. Dev Dyn. 237(3):527–544.

34. Hu Q, Duncan FE, Nowakowski AB, Antipova OA, Woodruff TK, O’Halloran TV, Wolfner MF. 2020. Zinc Dynamics during Drosophila Oocyte Maturation and Egg Activation. iScience. 23(7):101275.

35. Intra J, Cenni F, Perotti M-E. 2006. An alpha-L-fucosidase potentially involved in fertilization is present on Drosophila spermatozoa surface. Mol Reprod Dev. 73(9):1149–1158.

36. Kaur R, Leigh BA, Ritchie IT, Bordenstein SR. 2022. The Cif proteins from Wolbachia prophage WO modify sperm genome integrity to establish cytoplasmic incompatibility. PLoS Biol. 20(5):e3001584.

37. Kosakovsky Pond SL, Frost SDW. 2005. Not so different after all: a comparison of methods for detecting amino acid sites under selection. Mol Biol Evol. 22(5):1208–1222.

38. Kosakovsky Pond SL, Posada D, Gravenor MB, Woelk CH, Frost SDW. 2006. Automated phylogenetic detection of recombination using a genetic algorithm. Mol Biol Evol. 23(10):1891–1901.

39. Kracklauer MP, Wiora HM, Deery WJ, Chen X, Bolival B Jr, Romanowicz D, Simonette RA, Fuller MT, Fischer JA, Beckingham KM. 2010. The Drosophila SUN protein Spag4 cooperates with the coiled-coil protein Yuri Gagarin to maintain association of the basal body and spermatid nucleus. J Cell Sci. 123(Pt 16):2763–2772.

40. Kumar S, Suleski M, Craig JM, Kasprowicz AE, Sanderford M, Li M, Stecher G, Hedges SB. 2022. TimeTree 5: An Expanded Resource for Species Divergence Times. Mol Biol Evol. 39(8):msac174.

41. LaFlamme BA, Ram KR, Wolfner MF. 2012. The Drosophila melanogaster seminal fluid protease “seminase” regulates proteolytic and post-mating reproductive processes. PLoS Genet. 8(1):e1002435.

42. Lange A, Patel PH, Heames B, Damry AM, Saenger T, Jackson CJ, Findlay GD, Bornberg-Bauer E. 2021. Structural and functional characterization of a putative de novo gene in Drosophila. Nat Commun. 12(1):1667.

43. Lehti MS, Sironen A. 2016. Formation and function of the manchette and flagellum during spermatogenesis. J Reprod Fertil. 151(4):R43–54.

44. Levine MT, Jones CD, Kern AD, Lindfors HA, Begun DJ. 2006. Novel genes derived from noncoding DNA in Drosophila melanogaster are frequently X-linked and exhibit testis-biased expression. Proc Natl Acad Sci U S A. 103(26):9935–9939.

45. Lindsley DL, Tokuyasu KT. 1980. Spermatogenesis. In: Ashburner M, Wright TRF, editors. The Genetics and Biology of Drosophila, Volume 2D. London: Academic Press. p. 225–294.

46. Li P, Messina G, Lehner CF. 2023. Nuclear elongation during spermiogenesis depends on physical linkage of nuclear pore complexes to bundled microtubules by Drosophila Mst27D. PLoS Genet. 19(7):e1010837.

47. Long M, VanKuren NW, Chen S, Vibranovski MD. 2013. New gene evolution: little did we know. Annu Rev Genet. 47(1):307–333.

48. Loppin B, Berger F, Couble P. 2001. The Drosophila maternal gene sésame is required for sperm chromatin remodeling at fertilization. Chromosoma. 110(6):430–440.

49. Loppin B, Bonnefoy E, Anselme C, Laurençon A, Karr TL, Couble P. 2005. The histone H3.3 chaperone HIRA is essential for chromatin assembly in the male pronucleus. Nature. 437(7063):1386–1390.

50. Loppin B, Dubruille R, Horard B. 2015. The intimate genetics of Drosophila fertilization. Open Biol. 5(8):150076.

51. Loppin B, Lepetit D, Dorus S, Couble P, Karr TL. 2005. Origin and neofunctionalization of a Drosophila paternal effect gene essential for zygote viability. Curr Biol. 15(2):87–93.

52. Lüpold S, Manier MK, Puniamoorthy N, Schoff C, Starmer WT, Luepold SHB, Belote JM, Pitnick S. 2016. How sexual selection can drive the evolution of costly sperm ornamentation. Nature. 533(7604):535–538.

53. Manier MK, Belote JM, Berben KS, Novikov D, Stuart WT, Pitnick S. 2010. Resolving mechanisms of competitive fertilization success in Drosophila melanogaster. Science. 328(5976):354–357.

54. McLysaght A, Hurst L. 2016. Open questions in the study of de novo genes: what, how and why. Nat Rev Genet. 17:567–578.

55. Murrell B, Weaver S, Smith MD, Wertheim JO, Murrell S, Aylward A, Eren K, Pollner T, Martin DP, Smith DM, et al. 2015. Gene-wide identification of episodic selection. Mol Biol Evol. 32(5):1365–1371.

56. Ni J-Q, Zhou R, Czech B, Liu L-P, Holderbaum L, Yang-Zhou D, Shim H-S, Tao R, Handler D, Karpowicz P, et al. 2011. A genome-scale shRNA resource for transgenic RNAi in Drosophila. Nat Methods. 8(5):405–407.

57. Ødum MT, Teufel F, Thumuluri V, Almagro Armenteros JJ, Johansen AR, Winther O, Nielsen H. 2024. DeepLoc 2.1: multi-label membrane protein type prediction using protein language models. Nucleic Acids Res. 52(W1):W215–W220.

58. Ohsako T, Shirakami M, Oiwa K, Ibaraki K, Karr TL, Tomaru M, Sanuki R, Takano-Shimizu-Kouno T. 2021. The Drosophila Neprilysin 4 gene is essential for sperm function following sperm transfer to females. Genes Genet Syst. 96(4):177–186.

59. Ohsako T, Yamamoto M-T. 2011. Sperm of the wasted mutant are wasted when females utilize the stored sperm in Drosophila melanogaster. Genes Genet Syst. 86(2):97–108.

60. Okabe M. 2016. The Acrosome Reaction: A Historical Perspective. Adv Anat Embryol Cell Biol. 220:1–13.

61. Öztürk-Çolak A, Marygold SJ, Antonazzo G, Attrill H, Goutte-Gattat D, Jenkins VK, Matthews BB, Millburn G, Dos Santos G, Tabone CJ, et al. 2024. FlyBase: updates to the Drosophila genes and genomes database. Genetics. 227(1):iyad211.

62. Patlar B, Jayaswal V, Ranz JM, Civetta A. 2021. Nonadaptive molecular evolution of seminal fluid proteins in Drosophila. Evolution. 75(8):2102–2113.

63. Peng J, Zhao L. 2024. The origin and structural evolution of de novo genes in Drosophila. Nat Commun. 15(1):810.

64. Perotti ME, Cattaneo F, Pasini ME, Vernì F, Hackstein JH. 2001. Male sterile mutant casanova gives clues to mechanisms of sperm-egg interactions in Drosophila melanogaster. Mol Reprod Dev. 60(2):248–259.

65. Pitnick S, Hosken DJ, Birkhead TR. 2009. Sperm morphological diversity. In: Birkhead TR, Hosken DJ, Pitnick S, editors. Sperm Biology. London: Academic Press. p. 69–149.

66. Pitnick S, Markow T, Spicer GS. 1999. Evolution of multiple kinds of female sperm-storage organs in Drosophila. Evolution. 53(6):1804.

67. Ramm SA, Schärer L, Ehmcke J, Wistuba J. 2014. Sperm competition and the evolution of spermatogenesis. Molecular human reproduction. 20(12):1169–1179.

68. Rathke C, Baarends WM, Awe S, Renkawitz-Pohl R. 2014. Chromatin dynamics during spermiogenesis. Biochim Biophys Acta. 1839(3):155–168.

69. Ravi Ram K, Wolfner MF. 2007. Sustained post-mating response in Drosophila melanogaster requires multiple seminal fluid proteins. PLoS Genet. 3(12):e238.

70. Rele CP, Sandlin KM, Leung W, Reed LK. 2022. Manual annotation of Drosophila genes: a Genomics Education Partnership protocol. F1000Res. 11:1579.

71. Rivard EL, Ludwig AG, Patel PH, Grandchamp A, Arnold SE, Berger A, Scott EM, Kelly BJ, Mascha GC, Bornberg-Bauer E, et al. 2021. A putative de novo evolved gene required for spermatid chromatin condensation in Drosophila melanogaster. PLoS Genet. 17(9):e1009787.

72. Sakai A, Schwartz BE, Goldstein S, Ahmad K. 2009. Transcriptional and developmental functions of the H3.3 histone variant in Drosophila. Curr Biol. 19(21):1816–1820.

73. Santel A, Winhauer T, Blümer N, Renkawitz-Pohl R. 1997. The Drosophila don juan (dj) gene encodes a novel sperm specific protein component characterized by an unusual domain of a repetitive amino acid motif. Mech Dev. 64(1-2):19–30.

74. Schärer L, Da Lage J-L, Joly D. 2008. Evolution of testicular architecture in the Drosophilidae: a role for sperm length. BMC Evol Biol. 8:143.

75. Shetterly ML. 2016. Hidden Figures: The American Dream and the Untold Story of the Black Women Mathematicians Who Helped Win the Space Race. HarperCollins.

76. Soulsbury CD, Iossa G. 2024. Coevolution between eggs and sperm of insects. Proc Biol Sci. 291(2026):20240525.

77. Stromberg KA, Spain T, Tomlin SA, Powell J, Amarillo KD, Schroeder CM. 2023. Evolutionary diversification reveals distinct somatic versus germline cytoskeletal functions of the Arp2 branched actin nucleator protein. Curr Biol. 33(24):5326–5339.e7.

78. Suarez SS. 2008. Regulation of sperm storage and movement in the mammalian oviduct. Int J Dev Biol. 52(5-6):455–462.

79. Suvorov A, Kim BY, Wang J, Armstrong EE, Peede D, D’Agostino ERR, Price DK, Waddell P, Lang M, Courtier-Orgogozo V, et al. 2022. Widespread introgression across a phylogeny of 155 Drosophila genomes. Curr Biol. 32(1):111–123.e5.

80. Tautz D, Domazet-Lošo T. 2011. The evolutionary origin of orphan genes. Nat Rev Genet. 12(10):692–702.

81. Tirmarche S, Kimura S, Dubruille R, Horard B, Loppin B. 2016. Unlocking sperm chromatin at fertilization requires a dedicated egg thioredoxin in Drosophila. Nat Commun. 7:13539.

82. Vakirlis N, Acar O, Hsu B, Castilho Coelho N, Van Oss SB, Wacholder A, Medetgul-Ernar K, Bowman RW 2nd, Hines CP, Iannotta J, et al. 2020. De novo emergence of adaptive membrane proteins from thymine-rich genomic sequences. Nat Commun. 11(1):781.

83. Vakirlis N, Vance Z, Duggan KM, McLysaght A. 2022. De novo birth of functional microproteins in the human lineage. Cell Rep. 41(12):111808.

84. VanKuren NW, Long M. 2018. Gene duplicates resolving sexual conflict rapidly evolved essential gametogenesis functions. Nat Ecol Evol. 2(4):705–712.

85. Van Oss SB, Carvunis A-R. 2019. De novo gene birth. PLoS Genet. 15(5):e1008160.

86. Vedelek V, Bodai L, Grézal G, Kovács B, Boros IM, Laurinyecz B, Sinka R. 2018. Analysis of Drosophila melanogaster testis transcriptome. BMC Genomics. 19(1):697.

87. Wagstaff BJ, Begun DJ. 2005. Comparative genomics of accessory gland protein genes in Drosophila melanogaster and D. pseudoobscura. Mol Biol Evol. 22(4):818–832.

88. Weaver S, Shank SD, Spielman SJ, Li M, Muse SV, Kosakovsky Pond SL. 2018. Datamonkey 2.0: A Modern Web Application for Characterizing Selective and Other Evolutionary Processes. Mol Biol Evol. 35(3):773–777.

89. Wei KH-C, Chatla K, Bachtrog D. 2024. Single-cell RNA-seq of Drosophila miranda testis reveals the evolution and trajectory of germline sex chromosome regulation. PLoS Biol. 22(4):e3002605.

90. Weisman CM, Murray AW, Eddy SR. 2020. Many, but not all, lineage-specific genes can be explained by homology detection failure. PLoS Biol. 18(11):e3000862.

91. White-Cooper H, Doggett K, Ellis RE. 2009. The evolution of spermatogenesis. In: Birkhead TR, Hosken DJ, Pitnick S, editors. Sperm Biology. London: Academic Press. p. 151–183.

92. Wilburn DB, Swanson WJ. 2016. From molecules to mating: Rapid evolution and biochemical studies of reproductive proteins. J Proteomics. 135:12–25.

93. Wilson KL, Fitch KR, Bafus BT, Wakimoto BT. 2006. Sperm plasma membrane breakdown during Drosophila fertilization requires sneaky, an acrosomal membrane protein. Development. 133(24):4871–4879.

94. Wisotsky SR, Kosakovsky Pond SL, Shank SD, Muse SV. 2020. Synonymous Site-to-Site Substitution Rate Variation Dramatically Inflates False Positive Rates of Selection Analyses: Ignore at Your Own Peril. Mol Biol Evol. 37(8):2430–2439.

95. Wolfner MF, Suarez SS, Dorus S. 2023. Suspension of hostility: Positive interactions between spermatozoa and female reproductive tracts. Andrology. 11(5):943–947.

96. Yamaki T, Yasuda GK, Wakimoto BT. 2016. The Deadbeat Paternal Effect of Uncapped Sperm Telomeres on Cell Cycle Progression and Chromosome Behavior in Drosophila melanogaster. Genetics. 203(2):799–816.

97. Yang Z, Nielsen R, Goldman N, Pedersen AM. 2000. Codon-substitution models for heterogeneous selection pressure at amino acid sites. Genetics. 155(1):431–449.

98. Zhang L, Ren Y, Yang T, Li G, Chen J, Gschwend AR, Yu Y, Hou G, Zi J, Zhou R, et al. 2019. Rapid evolution of protein diversity by de novo origination in Oryza. Nature ecology & evolution. 3(4):679–690.

99. Zhao L, Saelao P, Jones CD, Begun DJ. 2014. Origin and spread of de novo genes in Drosophila melanogaster populations. Science. 343(6172):769–772.

100. Zhao L, Svetec N, Begun DJ. 2024. De Novo Genes. Annu Rev Genet. 58. doi:10.1146/annurev-genet-111523-102413.

101. Zhao Q, Zheng Y, Li Y, Shi L, Zhang J, Ma D, You M. 2024. An Orphan Gene Enhances Male Reproductive Success in Plutella xylostella. Mol Biol Evol. 41(7). doi:10.1093/molbev/msae142.

