## Supplementary material for "An orphan gene is essential for efficient sperm entry into eggs in *Drosophila melanogaster*": Figure S1

## A)

Oligos annealed to create the pValium20 insert for targeting *CG43167* (*kj*) expression:

CG43167-top: CTAGCAGT**CGGCAGCTAAAGTATTGCGAA**TAGTTATATTCAAGCATA**TTCGCAATACTTTAGCTGCC**GGCG
CG43167-bot: AATTCGC**CGGCAGCTAAAGTATTGCGAA**TATGCTTGAATATAACTA**TTCGCAATACTTTAGCTGCC**GACTG

Coordinates of targeted sequence in *CG43167* gene: chr2L:11,657,752-11,657,77

## B)

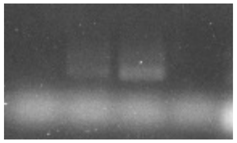


## C)

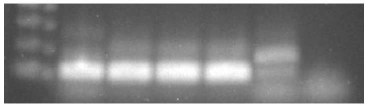


*indicates cDNA samples from flies knocked down for a different candidate *de novo* gene

## D)

RT-PCR primers:

CG43167-F: CTTTATTTCCGTGGCGTTTC
CG43167-R: TTTTCTTTCGTCTCGGTTCG
