## Supplementary material for "An orphan gene is essential for efficient sperm entry into eggs in *Drosophila melanogaster*": Figure S2

### A)

5’ gRNA target site = chr2L: 11657428-11657403
3’ gRNA target site = chr2L: 11658105-11658080

Deleted region: chr2L: 11657407-11658095

Primers for screening for the deletion:
CG43167-screen-F: GCCAAAACTGAATCCTTGAGAGC
CG43167-screen-R: GTCAACTGCCCCTTTTCTGG

### B)

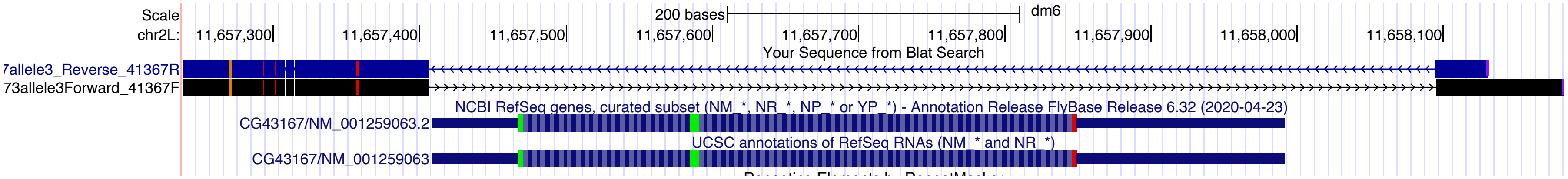
