## Supplementary figures and images for "An orphan gene is essential for efficient sperm entry into eggs in *Drosophila melanogaster*"

### Figure S3

**Figure S3**

**A**

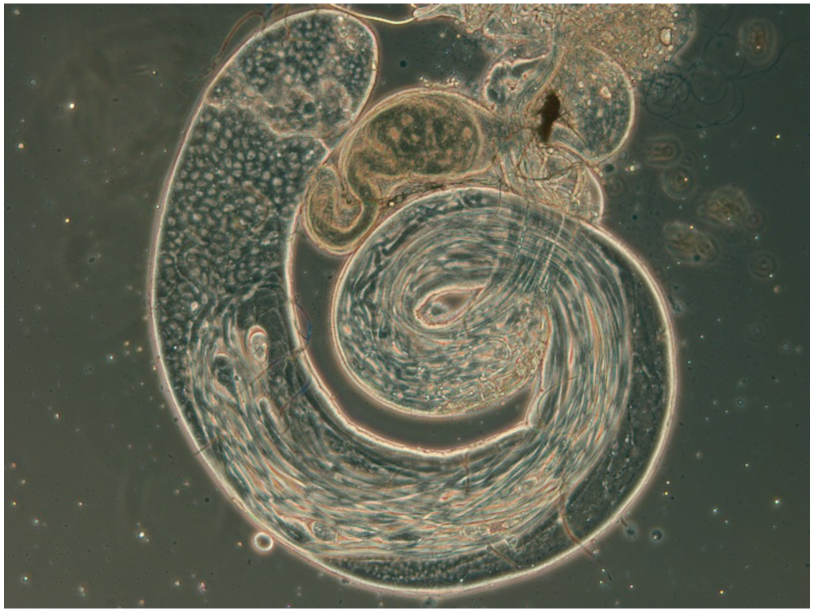

*kj⁺/kj⁺*

**B**

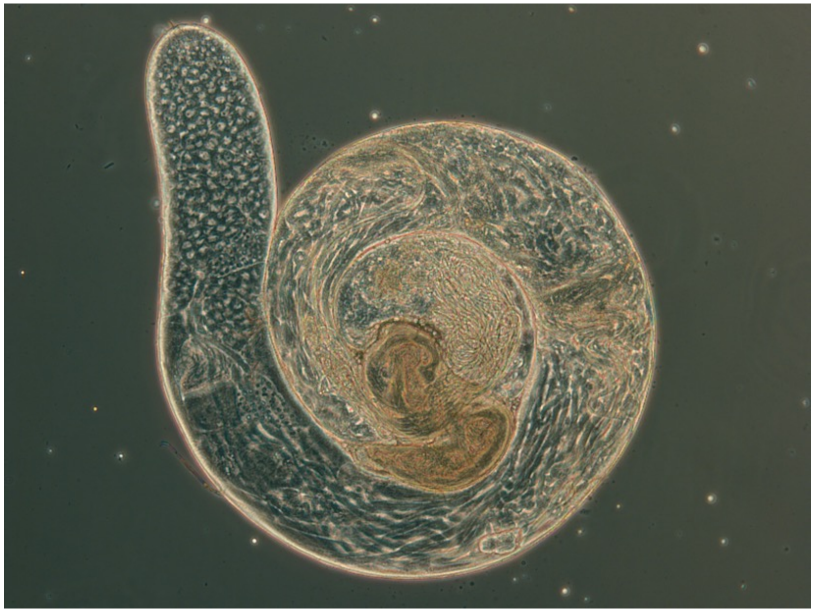

*Δkj/Δkj*

### Figure S4

**Figure S4**

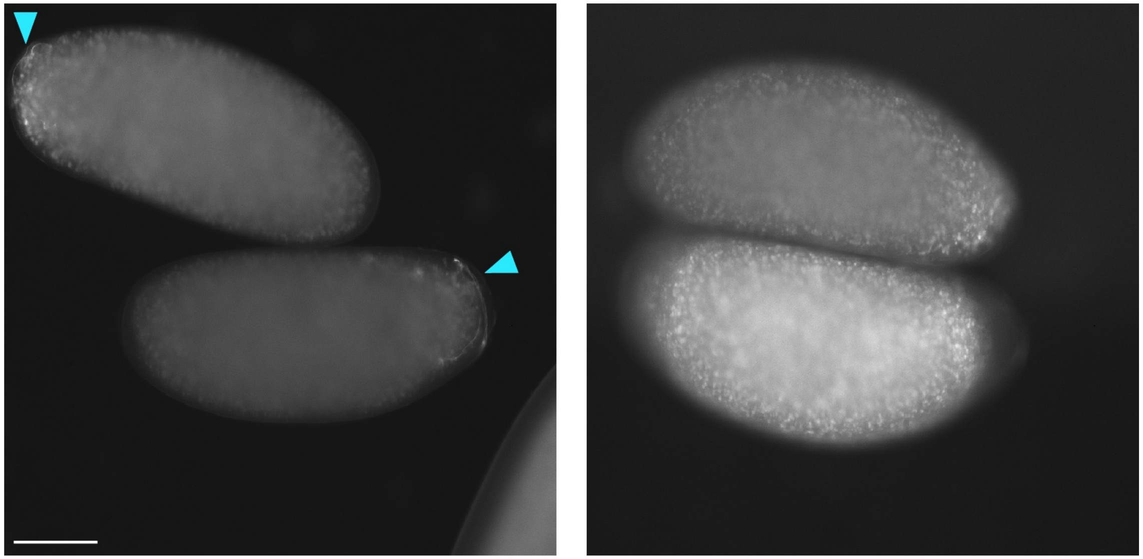

♂Δ*kj*/CyO, ♀WT　　♂Δ*kj*/Δ*kj*, ♀WT

### Figure S5

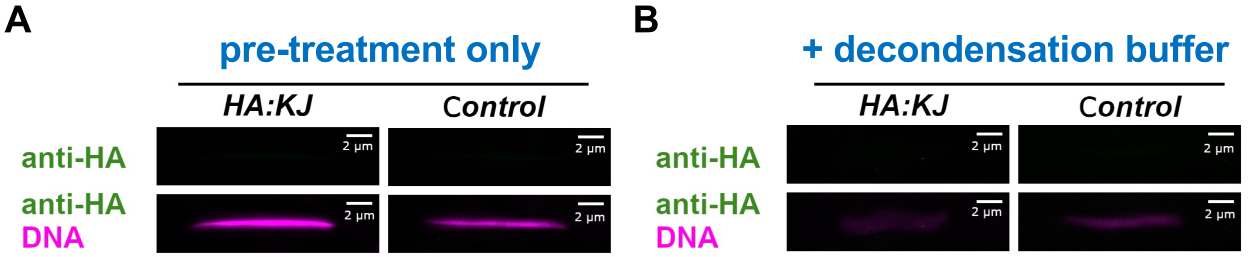

### Figure S6

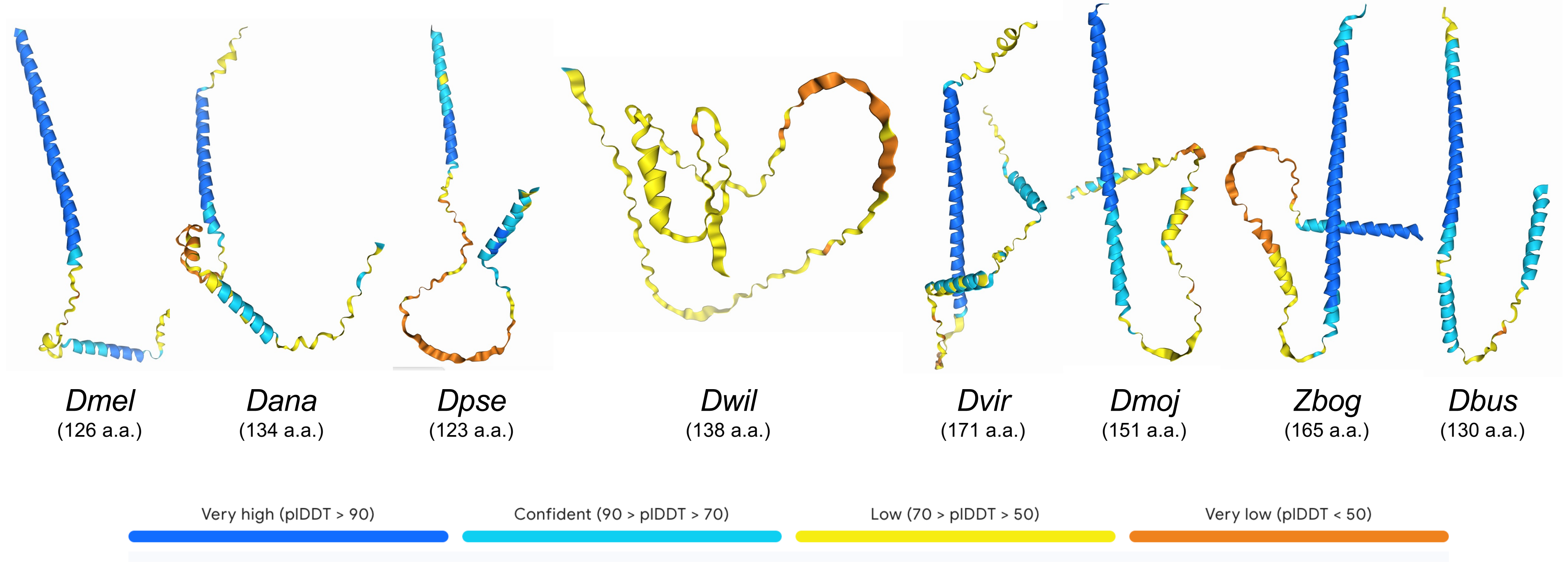
