## Supplementary material for "An orphan gene is essential for efficient sperm entry into eggs in *Drosophila melanogaster*": Figure S7

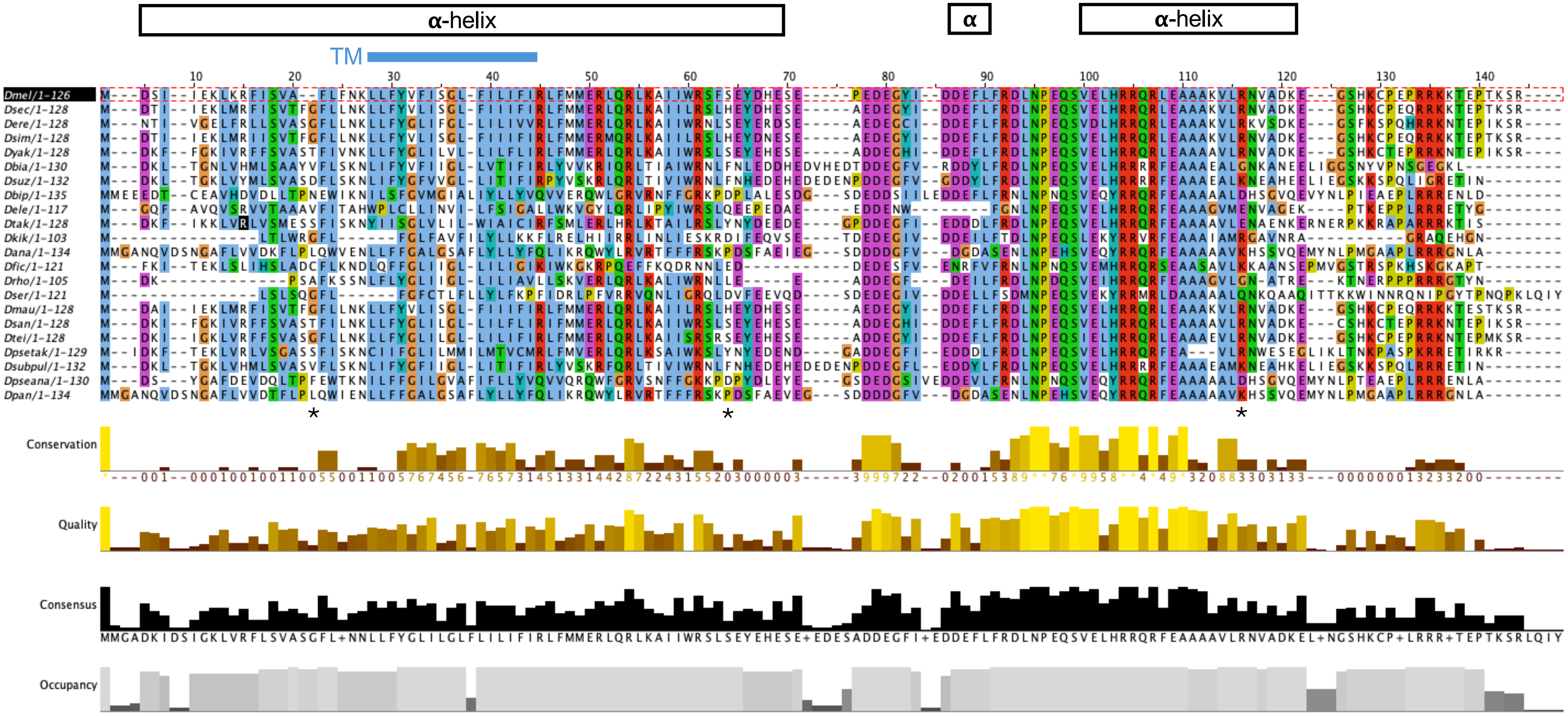


For the *mel* ortholog, the positions of the AlphaFold3-predicted alpha helices are indicated by black boxes, and the position of the DeepTMHMM-predicted transmembrane (TM) domain is shown by the blue line. * indicates alignment positions identified by FEL as have evolved under recurrent diversifying (positive) selection.
