## Supplementary material for "An orphan gene is essential for efficient sperm entry into eggs in *Drosophila melanogaster*": Figure S8

### Figure S6

**A)** *D. melanogaster* genome:

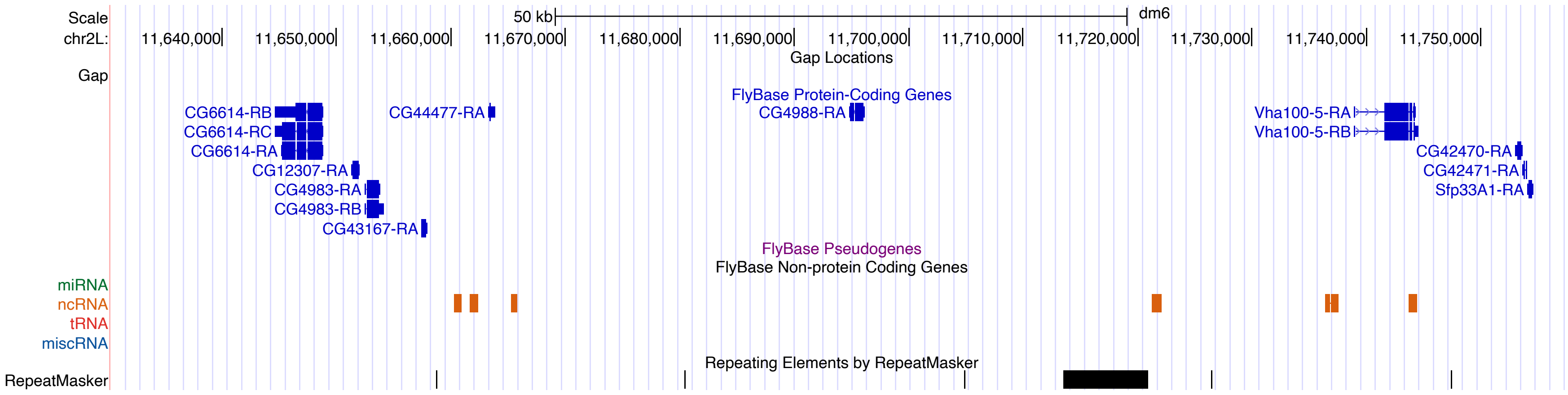


CG43167 (*kj*) and surrounding genes on chr 2L of *D. melanogaster*.

Upstream genes include ***CG6614***, *CG12307* and ***CG4983***.
Downstream gnes include *CG44477*, *CG4988* and ***Vha100-5***.

Genes in bold are in conserved positions around the putative *kj* ortholog in *D. virilis*.

**B)** *D. virilis* genome:

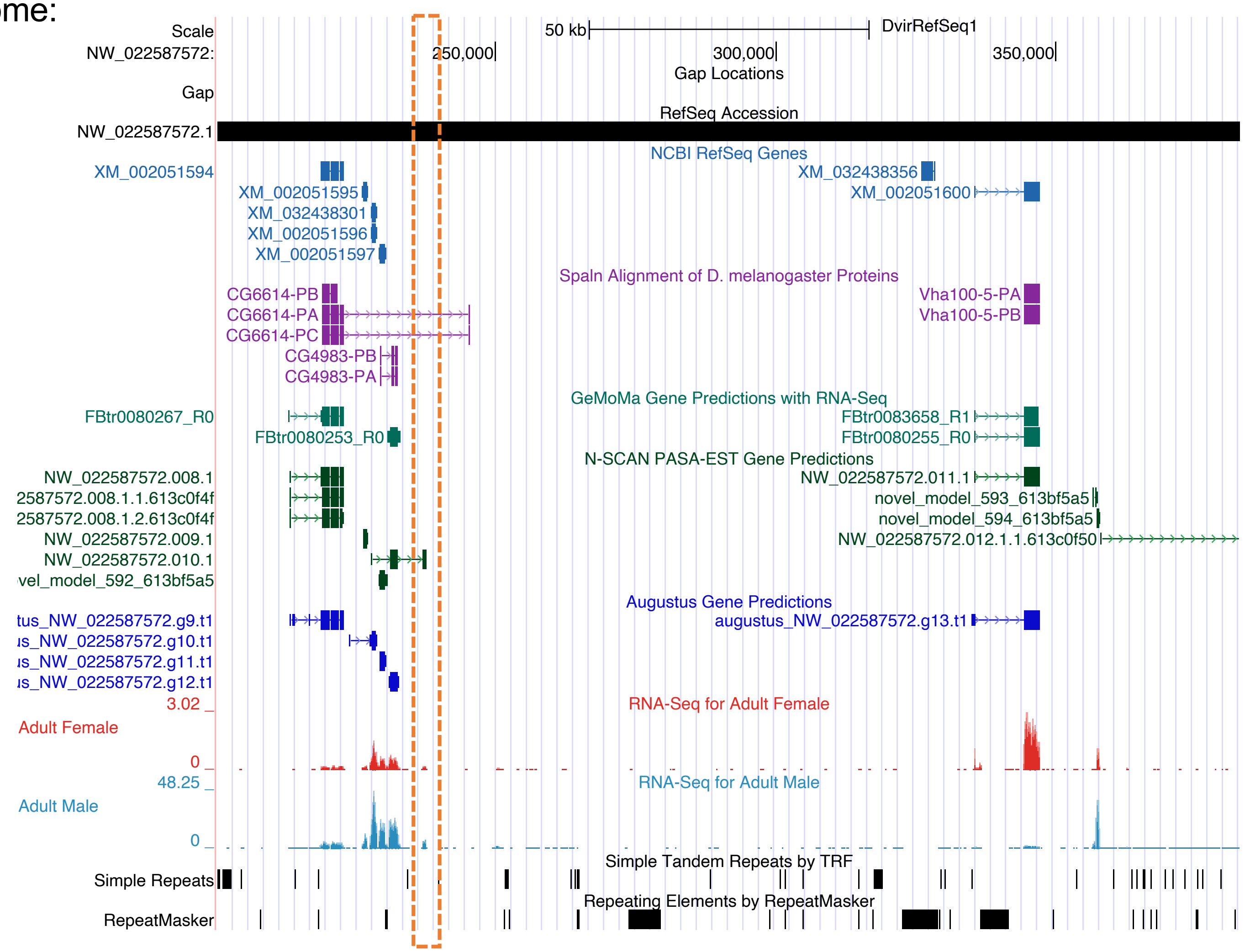

### C) *D. virilis* genome:

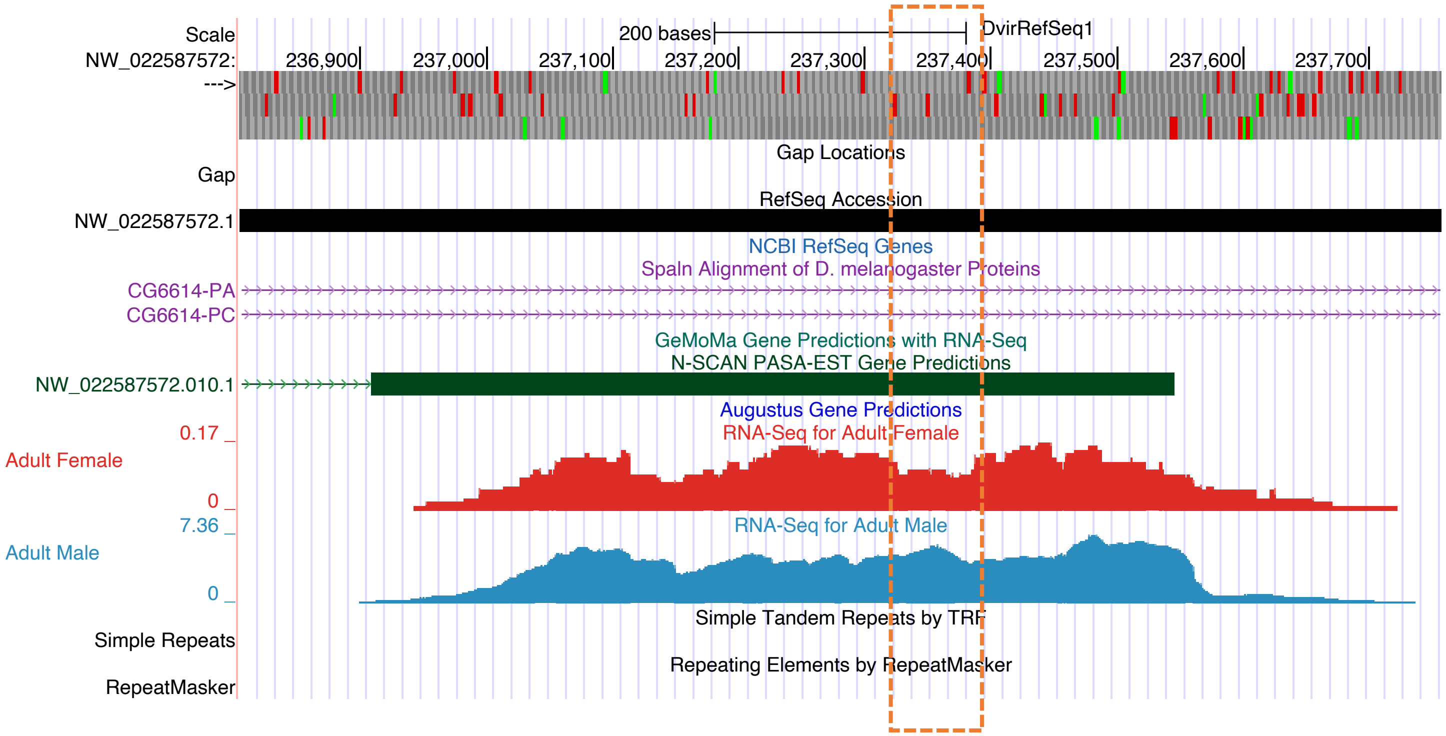


```
TBLASTN hit:
mel  Query 76          FRDLNPEQSVELHRRQRLEAAAKVL 100
                       FR LNP QSVE +R+R E+ A ++
vir  Sbjct 237321      FRRLNPIQSVERQQRKRSESVAAIV 237395
(frame +3)
```

**D)** *D. virilis* ORF surrounding TBLASTN hit*:*

**MLNTTSILSKMPLLQSLEISDNAHLFIAVGIALCILLWYAAKFESDQILMDVLRKRQQQLQELQRKHQLKLLLKKTTKSE PSAPSTEESLTPQANHTFRRLNPIQSVERQQRKRSESVAAIVAAGDASATPRLRTNFKCLKPDQSVERHRRMRANLKMNV PDFAETKQKTD***

BLASTP: *D. virilis* ORF (query) vs. *D. melanogaster* KJ (subject):

**Dmel_KJ**

**Sequence ID:** **Query_94983** **Length:** **126** **Number of Matches:** **2**

**Range 1: 11 to 100** Graphics ▼ Next Match ▲ Previous Match

| Score | Expect | Method | Identities | Positives | Gaps |
|---|---|---|---|---|---|
| 34.7 bits(78) | 1e-07 | Compositional matrix adjust. | 27/97(28%) | 45/97(46%) | 7/97(7%) |

```
Query  26   FIAVGIALCILLWYAAKFESDQILMDVLRKRQQQLQELQRKHQLKLLLKKTTKSEPSAPS  85
            FI+V      LL+Y   F S   ++  +R     L  ++R  +LK ++ ++
Sbjct  11   FISVAFLFNKLLFYV--FISGLFILIFIR-----LFMMERLQRLKAIIWRSFSEYDHESE  63

Query  86   TEESLTPQANHTFRRLNPIQSVERQQRKRSESVAAIV  122
             E+         FR LNP QSVE  +R+R E+ A ++
Sbjct  64   PEDEGYIDDEFLFRDLNPEQSVELHRRQRLEAAAKVL  100
```

**Range 2: 76 to 105** Graphics ▼ Next Match ▲ Previous Match ▲ First Match

| Score | Expect | Method | Identities | Positives | Gaps |
|---|---|---|---|---|---|
| 26.6 bits(57) | 6e-05 | Compositional matrix adjust. | 15/30(50%) | 18/30(60%) | 4/30(13%) |

```
Query  137  FKCLKPDQSVERHRRMR----ANLKMNVPD  162
            F+ L P+QSVE HRR R    A +  NV D
Sbjct  76   FRDLNPEQSVELHRRQRLEAAAKVLRNVAD  105
```

**E)** DeepTMHMM topology predictions:

*Dmel* KJ:

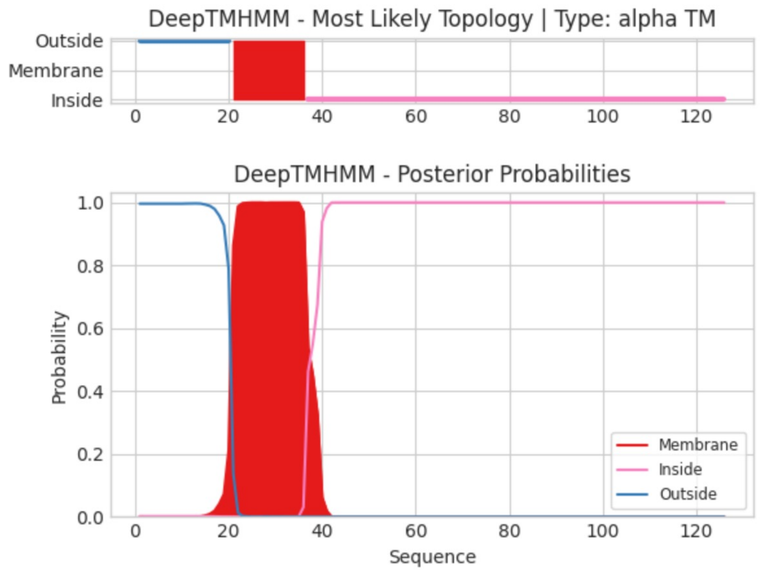


*Dvir* ORF:

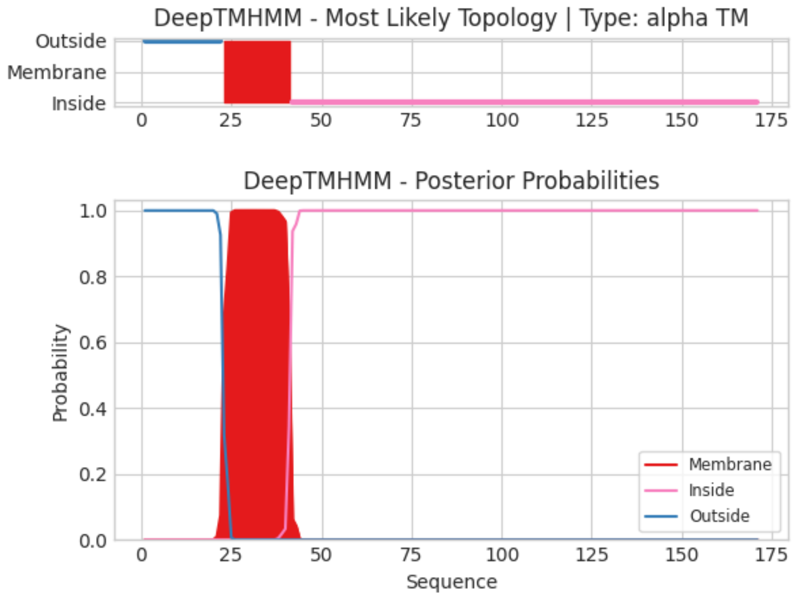
