## Supplementary material for "An orphan gene is essential for efficient sperm entry into eggs in *Drosophila melanogaster*": Figure S9

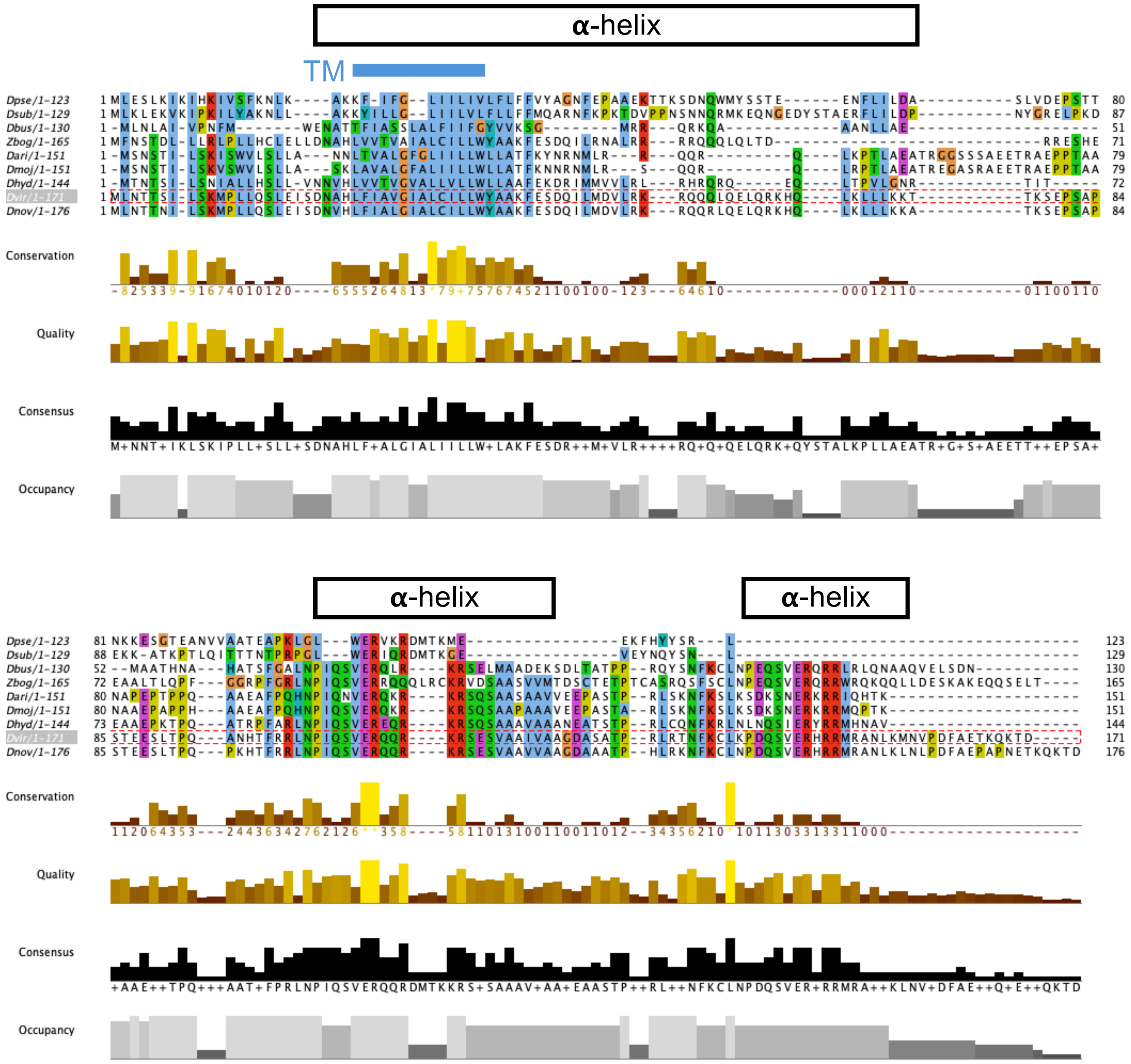


The positions of the AlphaFold3-predicted alpha helices (boxes) and the DeepTMHMM-predicted transmembrane (TM) domain (blue line) are indicated for the *D. virilis* ortholog.
