## Supplementary material for "An orphan gene is essential for efficient sperm entry into eggs in *Drosophila melanogaster*": Figure S10

### Figure S7

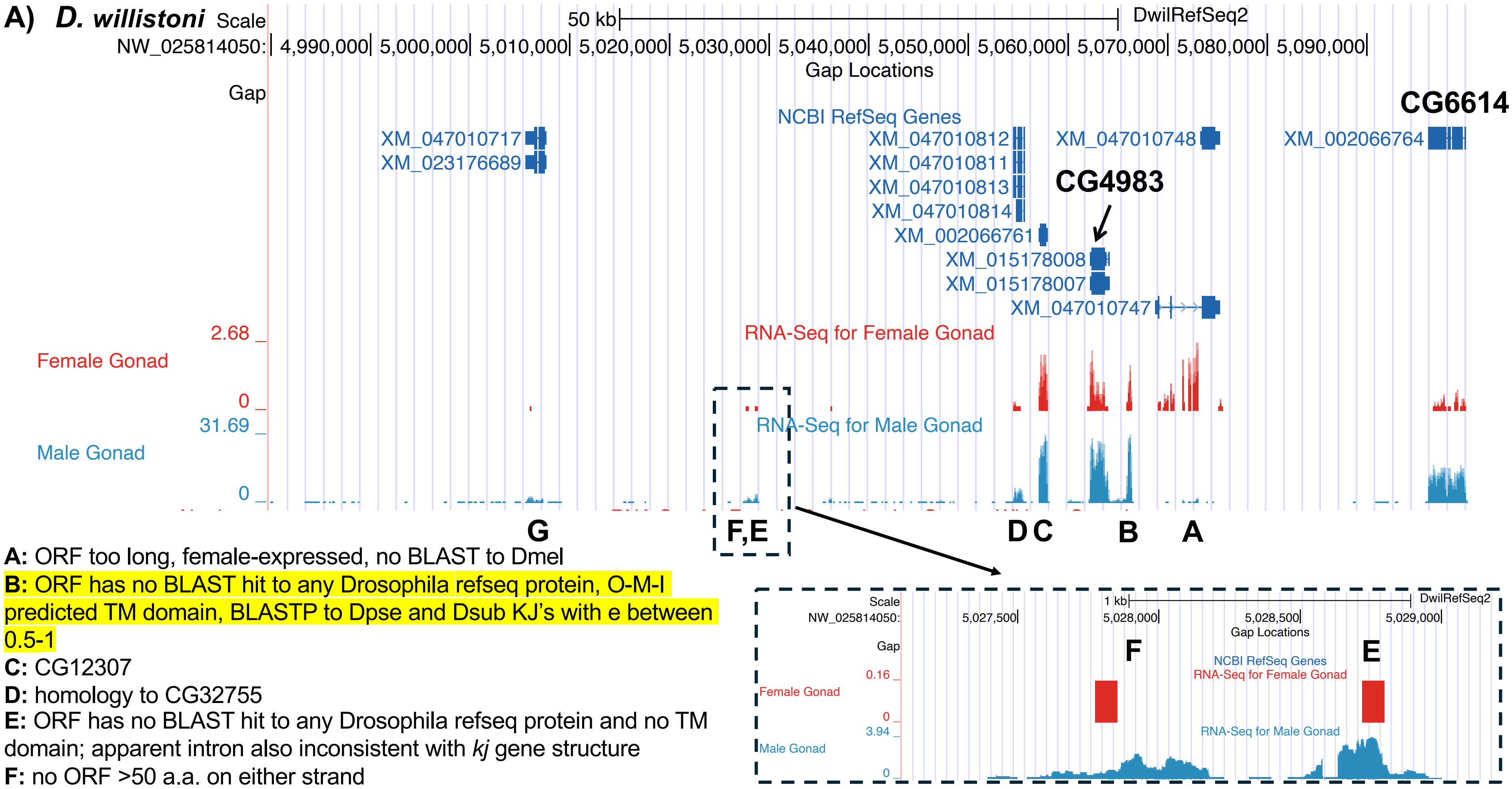


**A:** ORF too long, female-expressed, no BLAST to Dmel

**B:** ORF has no BLAST hit to any Drosophila refseq protein, O-M-I predicted TM domain, BLASTP to Dpse and Dsub KJ's with e between 0.5-1

**C:** CG12307

**D:** homology to CG32755

**E:** ORF has no BLAST hit to any Drosophila refseq protein and no TM domain; apparent intron also inconsistent with *kj* gene structure

**F:** no ORF >50 a.a. on either strand

**G:** homology to CG32755

### B) ***D. willistoni*** possible ORF

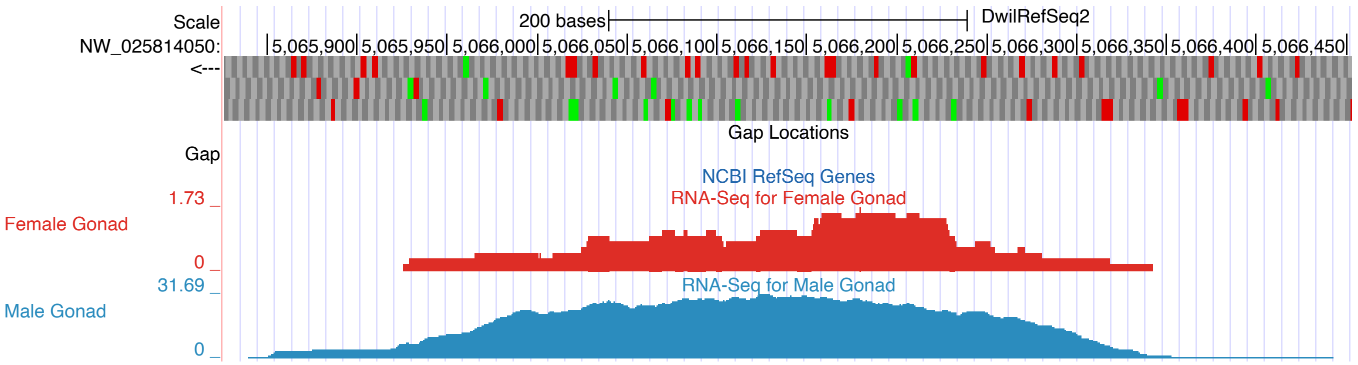


- No BLASTP similarity to any *Drosophila* RefSeq protein with e < 5.
- Pairwise BLASTP to potential KJ orthologs gives 0.5 < e < 1 for *Dpse* and *Dsub* (see next slide).
- Predicted TM domain with O-M-I topology.

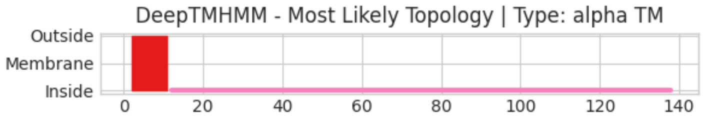


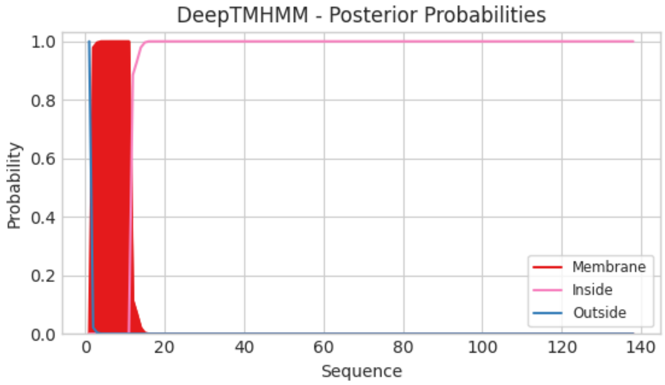

### C) *D. willistoni* possible ORF

The same region of the Dwil ORF has faint homology to different portions of the Dsub ORF.

Download Graphics Sort by: E value

**Dsub**

Sequence ID: **Query_2337022** Length: **129** Number of Matches: **2**

**Range 1: 59 to 90** Graphics ▼ Next Match ▲ Previous Match

| Score | Expect | Method | Identities | Positives | Gaps |
|---|---|---|---|---|---|
| 16.9 bits(32) | 0.53 | Compositional matrix adjust. | 13/33(39%) | 19/33(57%) | 1/33(3%) |

```
Query  53   IEEENLEDFNEGGEPHKQLDKATGTEDDKDKKK  85
            ++E+N ED++    E     LD    G E  KD+KK
Sbjct  59   MKEQNGEDYST-AERFLILDPNYGRELPKDEKK  90
```

**Range 2: 53 to 98** Graphics ▼ Next Match ▲ Previous Match ▲ First Match

| Score | Expect | Method | Identities | Positives | Gaps |
|---|---|---|---|---|---|
| 13.5 bits(23) | 8.2 | Compositional matrix adjust. | 11/46(24%) | 21/46(45%) | 6/46(13%) |

```
Query  86   NENDKDVKEMDEDTLSMEEN------NDGNDDREDETLLATPQVEL  125
            N N++ +KE + +  S  E        N G +  +DE      P +++
Sbjct  53   NSNNQRMKEQNGEDYSTAERFLILDPNYGRELPKDEKKATKPTLQI  98
```

Download Graphics

**Dpse (possible)**

Sequence ID: **Query_2337021** Length: **123** Number of Matches: **1**

**Range 1: 29 to 115** Graphics ▼ Next Match ▲ Previous Match

| Score | Expect | Method | Identities | Positives | Gaps |
|---|---|---|---|---|---|
| 16.2 bits(30) | 0.89 | Compositional matrix adjust. | 18/97(19%) | 43/97(44%) | 10/97(10%) |

```
Query  1    MLSIVFVYFVIRFSGKSKQIEKKPVADDPGSRIDEFEQYAEESDLDECDGTTIEEENLED  60
            +L ++F++FV   ++G  +       P A+       +++   + E +      D + ++E +  +
Sbjct  29   ILIVLFLFFV--YAGNFE-----PAAEKTTKSDNQWMYSSTEENFLILDASLVDEPSTTN  81

Query  61   FNEGGEPHKQLDKATGTEDDKDKKKNENDKDVKEMDE  97
              E G     + +     TE  K          +D+ +M+E
Sbjct  82   KKESGT---EANVVAATEAPKLGLWERVKRDMTKMEE  115
```
