## Supplementary material for "An orphan gene is essential for efficient sperm entry into eggs in *Drosophila melanogaster*": Figure S11

### Figure S8

#### A) *D. albomicans* syntenic region

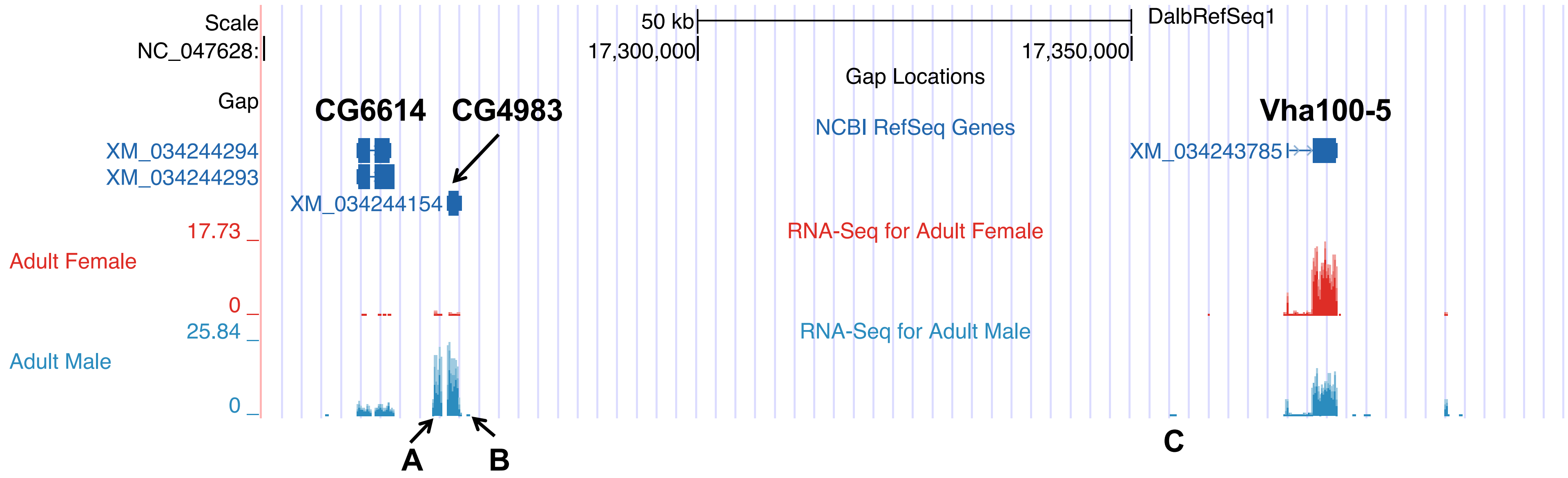


A: ORF interrupted by a fragment of a Jockey transposon; no part of the region has any BLASTX similarity with e < 10 to any non-mel group KJ ortholog
B: no ORFs > 30 a.a.
C: one ORF of 63 a.a., no TM domain, some parts of ORF have no RNAseq coverage

### B) *D. albomicans*, cont.

TBLASTN all non-mel group KJ orthologs against the syntenic region (150,000 bp) of the *D. albomicans* genome with an e = 10 cutoff. Nine of ten orthologs had no significant hit. The Dvir ortholog had an e = 0.69 hit to a region with no predicted ORF and no RNA-seq data. The e-value suggests this is probably a spurious hit, but if it were real, there is no evidence of an ORF in the identified reading frame or of transcription.

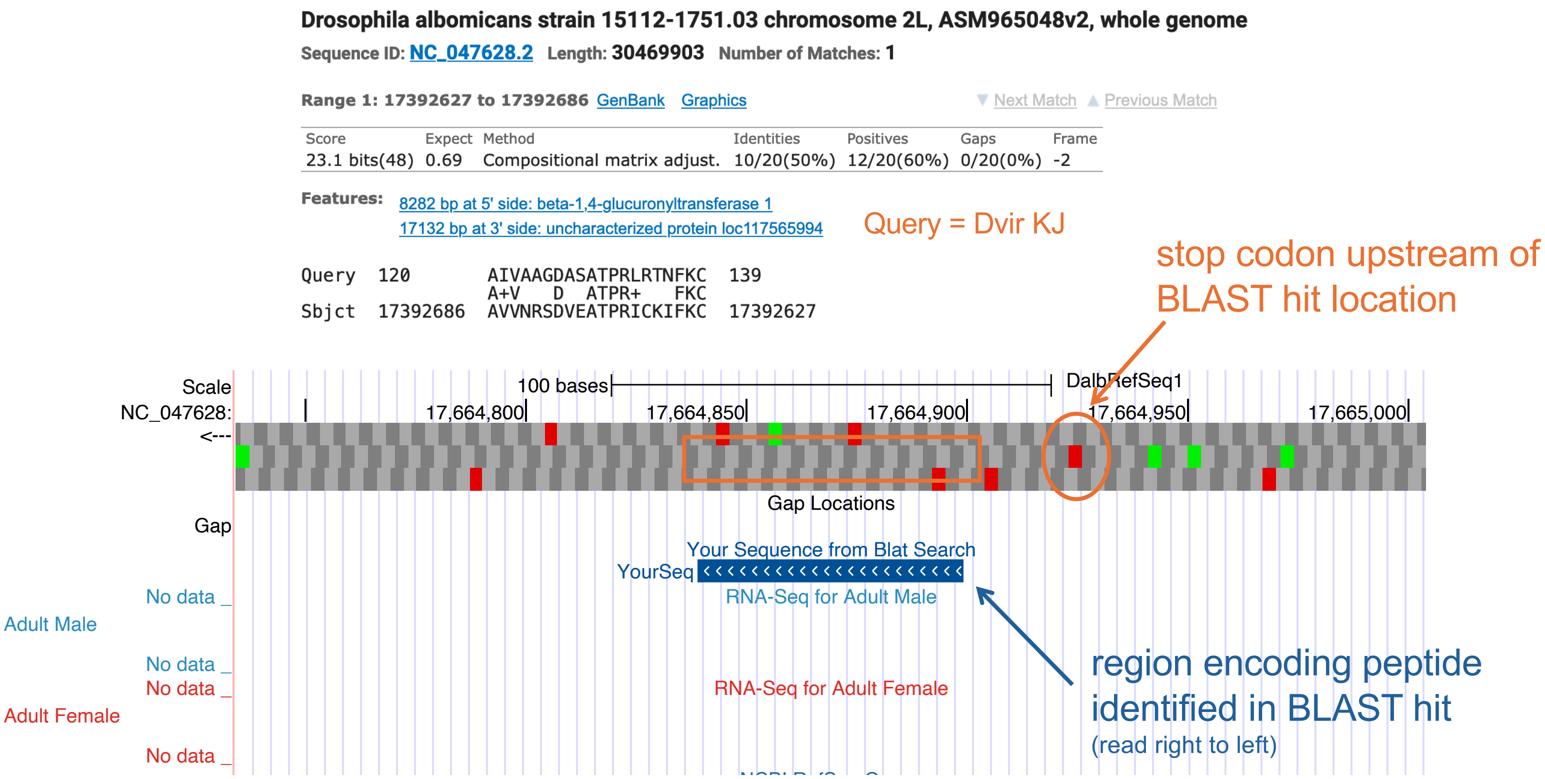

**C)** ***D. grimshawi***

TBLASTN search of *Dvir* KJ to the *Dgri* syntenic region found a hit to this location:

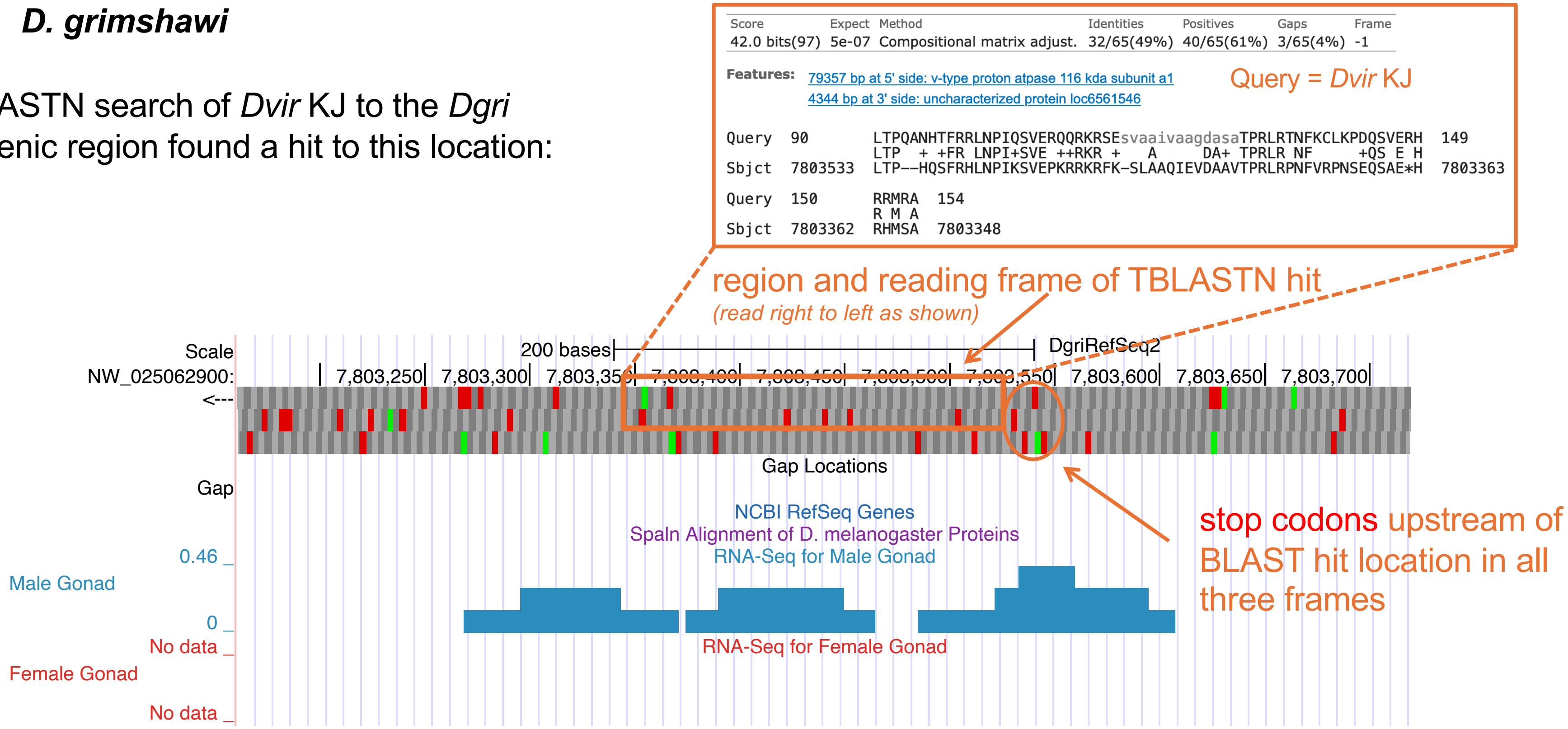


*Dvir* KJ amino acid positions 90-154 have a TBLASTN hit to the region boxed in orange. However, there are stop codons upstream (to the right in the diagram), suggesting there is no full-length ORF here. The RNAseq signal is also lower and spottier across the region than seen for other orthologs. These data suggest possible pseudogenization in *Dgri*, concomitant with a reduction in expression levels.
