## Supplementary material for "An orphan gene is essential for efficient sperm entry into eggs in *Drosophila melanogaster*": Table S1

**Table S1. Coordinates, protein lengths, predicted protein topologies, and expression data of putative *kj* orthologs in *melanogaster* group species.**

| Species | Accession | Length | DeepTMHMM topology prediction* | Location of CDS (for species with RNA-seq data available) | Adult expression (based on RNAseq data) |
|---|---|---|---|---|---|
| *D. melanogaster* | NP_001245992.1 | 126 a.a. | One: O-M-I | chr2L:11657469-11657849 | male-specific |
| *sechellia* | manual annotation based on XP_032578684.1 | 128 | One: O-M-I | NC_045949.1:11411740-11412123 | male-specific |
| *erecta* | XP_001969822.1 | 128 | One: O-M-I | NW_020825200.1: 14045265-14045648 | male-expressed; no adult female data available |
| *simulans* | XP_016024676.1 | 128 | One: O-M-I | NC_052520.2:11546635-11547018 | strongly male-biased |
| *mauritiana* | XP_033173237.1 | 128 | One: O-M-I | | |
| *yakuba* | XP_002088383.2 | 128 | One: I-M-O | NC_052527.2:8161458-8161841 | male-specific |
| *santomea* | XP_039502387.1 | 128 | One: I-M-O | | |
| *teissieri* | XP_043662272.1 | 128 | no TM predicted | | |
| *biarmipes* | manual annotation based on XP_016963382.2 | 130 | One: O-M-I | NW_025319395.1:20535887-20536276 | male-specific |
| *pseudotakahashii* | KAH8353071.1 | 129 | no TM predicted | | |
| *subpulchrella* | XP_037711180.1 | 132 | One: O-M-I | | |
| *suzukii* | XP_016933652.1 | 132 | One: O-M-I | NW_023496800.1:3607824-3608219 | male-specific |
| *pseudoananassae* | KAH8317672.1 | 130 | One: O-M-I | | |
| *bipectinata* | XP_017098007.2 | 135 | One: O-M-I | NW_025063917.1:1971100-1971504 | male-specific |

| | | | | | |
|---|---|---|---|---|---|
| *elegans* | XP_041566325.1 | 117 | One: signal peptide-O-M-I | NW_024546167.1:4865698-4866048 | male-specific; possible untranslated first exon based on RNAseq data |
| *takahashii* | XP_017011948.2 | 128 | One: O-M-I | NW_025323500.1:19008812-19009195 | male-specific |
| *jambulina* | KAH8281563.1 | n/a | n/a | *only partial sequence (GenBank protein begins with L rather than M; excluded from further analysis) | |
| *birchii* | KAH8249223.1 | n/a | n/a | *only partial sequence (GenBank protein begins with K rather than M; excluded from further analysis) | |
| *kikkawai* | XP_017032871.1 | 103 | One: O-M-I | NW_024571435.1:21267242-21267550 | male-specific |
| *pandora* | KAH8325818.1 | 134 | One: O-M-I | | |
| *ananassae* | XP_001962181.1 | 134 | One: O-M-I | NC_057930.1:21962814-21963215 | male-biased |
| *setifemur* | KAH8412767.1 | 149 | no TM predicted | *broadly similar pattern of amino acids, but longer length made it difficult to align, so excluded from PAML analysis | |
| *ficusphila* | XP_017062242.1 | 121 | One: O-M-I | NW_025064537.1:1800760-1801122 | male-specific |
| *bunnanda* | KAH8242010.1 | n/a | n/a | *only partial sequence (GenBank protein begins with T rather than M; excluded from further analysis) | |
| *rhopaloa* | XP_016980774.1 | 105 | One: O-M-I | NW_025335083.1:5929151-5929465 | male-specific |
| *serrata* | manual annotation based on KAH8356241.1 | 121 | One: O-M-I | NW_018367237.1:578531-578169 | male-specific |
| *gunungcola* | KAI8043326.1 | 68 | no TM predicted | *only aligns to the C-terminal portion of the protein; may be only a partial sequence, excluded from PAML analysis | |
| *eugracilis*** | manual annotation based on XP_017065384.2 (error in genome assembly) | 119 | One: I-M-O | NW_024573311.1:3198974-3198617, excluding the T at position 3198827 (since this nucleotide is not present in most RNA-seq reads of this locus) | male-specific |

*DeepTMHMM structural predictions are listed from N-terminus to C-terminus. O-M-I indicates a predicted topology with the N terminus outside the membrane, a membrane-spanning domain, and the C terminus inside the membrane. I-M-O indicates a prediction of the N terminus inside the membrane, a membrane-spanning domain, and the C terminus outside the membrane. In one instance, a secretion signal peptide sequence was predicted at the N-terminus.
**Excluded from molecular evolutionary analyses for positive selection due to likely error in reference genome assembly.
