## Supplementary material for "An orphan gene is essential for efficient sperm entry into eggs in *Drosophila melanogaster*": Table S2

**Table S2. Coordinates, protein lengths, predicted protein topologies, and expression data of putative *kj* orthologs outside of the *melanogaster* group.**

| Species | Location/accession of CDS (no stop) | Length | DeepTMHMM topology prediction* | Adult expression (based on RNAseq data) |
|---|---|---|---|---|
| *D. virilis* | NW_022587572:237,030-237,524 (alternative start at 237,060) | 171 a.a. (or 161) | O-M-I | male-biased |
| *Z. bogoriensis* | KAH8418808.1 | 165 | O-M-I | n/a |
| *D. busckii* | NC_046604:14,102,436-14,102,047 | 130 | O-M-I | male-specific |
| *D. pseudoobscura* | NC_046681:20,365,368-20,365,000 | 123 | I-M-O | male-biased |
| *D. subobscura* | NC_048534.1:10,774,141-10,774,527 | 129 | I-M-O | expressed (only data available is whole body from pooled sexes) |
| *D. willistoni* | NW_025814050: 5066348-5065927 | 138 | O-M-I (only the M is O) | male-biased |
| *D. mojavensis* | NW_025318872.1:414,803-415,255 | 151 | O-M-I | male-specific |
| *D. arizonae* | NW_017127683.1:23,724,054-23,724,506 | 151 | O-M-I | n/a |
| *D. novamexicana* | NW_020824787.1:5265510-5264983 | 176 | O-M-I | male-specific, highly testis-enriched |
| *D. hydei* | NW_022045827.1:4521201-4521632 | 144 | O-M-I | male-specific |

*DeepTMHMM structural predictions are listed from N-terminus to C-terminus. O-M-I indicates a predicted topology with the N terminus outside the membrane, a membrane-spanning domain, and the C terminus inside the membrane. I-M-O indicates a prediction of the N terminus inside the membrane, a membrane-spanning domain, and the C terminus outside the membrane.
